# Phages infecting the common gut commensal *Escherichia coli* HS reveal tropism for its *Klebsiella*-like capsule

**DOI:** 10.64898/2026.05.19.726188

**Authors:** Aleksandr Shenfeld, Oksana Kotovskayga, Kirill Petrikov, Alla Golomidova, Alina Demkina, Maria Zavialova, Olha Dorozh, Olga Komarova, Polina Iarema, Valentina M. Krasilnikova, Nikolay V. Volozhantsev, Konstantin Severinov, Andrei Letarov, Artem Isaev

## Abstract

Tailed dsDNA phages are ubiquitous and thought to infect almost any bacterial species. *Escherichia coli* HS – a commensal strain used as a model in gut colonization studies – has not been reported as a host for dsDNA phages and is resistant to more than 100 coliphages from the BASEL collection. Here, we report the first phages infecting *E. coli* HS, characterize their interaction with an endogenous BREX system, and provide a detailed genomic and phylogenetic description. Phages ϕHS1 (*Queuovirinae*) and ϕHS2 (*Ackermannviridae*) possess dPreQ₀ and 5-*NeO*mdU modifications that confer resistance to restriction digestion. ϕHS3 is a temperate phage, related to the native HS prophage ϕHS4, and represents a founding member of a novel genus (*Hueyvirus*) of P22-like phages capable of lateral transduction. We reveal exchange of tailspike genes between ϕHS2, ϕHS3, and *Klebsiella*-specific phages. Furthermore, we demonstrate that *E. coli* HS encodes a *Klebsiella*-like K47 capsule required for phage infection. These results suggest that the broad phage resistance of *E. coli* HS may be linked to its capsular type.

## Introduction

*Escherichia coli* is one of the most common bacterial species of the healthy human gut microbiome (Tenaillon *et al*, 2010; Foster-Nyarko & Pallen, 2022) and, at the same time, one of the most concerning human pathogens (Tenaillon *et al*, 2010; Croxen & Finlay, 2010). The genetic features that distinguish pathogenic *E. coli* lineages from commensal strains, which replicate in the human gut without causing disease, remain challenging to identify (Rasko *et al*, 2008). Commensal bacteria establish a complex interaction with the host immune system and are considered beneficial, as they serve as a barrier against secondary colonization by pathogenic microorganisms (Leshem *et al*, 2020). *E. coli* HS, first isolated in 1958, is a human gut commensal strain, tolerated in doses up to 10¹⁰ cells (DuPont *et al*, 1971; Formal *et al*, 1958; Levine *et al*, 1978; Doranga *et al*, 2024). Pre-colonization of the mouse gut with *E. coli* HS inhibits establishment of the pathogenic population of *E. coli* O157:H7, likely through nutritional competition, while still permits colonization by the commensal *E. coli* Nissle 1917, which occupies a distinct nutritional niche (Leatham *et al*, 2009; Maltby *et al*, 2013).

To persist in the human gut, bacteria must evade host immune recognition and resist bacteriophage infection. Both processes are largely governed by bacterial surface structures, although bacteria also encode diverse intracellular antiviral defense systems (Georjon & Bernheim, 2023; Zheng *et al*, 2020). The outer surface of *E. coli* is composed of the O-antigen extensions of the membrane lipopolysaccharide, with more than 150 serotypes reported to date, as well as capsular polysaccharides, which are broadly classified into four groups (Rendueles *et al*, 2017; Liu *et al*, 2020a). Although a recent bioinformatic census revealed previously unappreciated diversity among *E. coli* type 2 and type 3 capsules (Gladstone *et al*, 2026; Miravet-Verde *et al*, 2026), type 1 and type 4 capsules remain significantly less understood. O-antigens and capsules can act as physical barriers to phage adsorption; conversely, some phages use these structures as receptors or degrade capsular polysaccharides via depolymerases (Knecht *et al*, 2020; Letarov, 2023; Leprince *et al*, 2026). Notably, *E. coli* HS has not been reported to be infected by tailed dsDNA phages, despite the prevalence of such phages in the human gut. In contrast, an F^+^ derivative of HS is susceptible to the F-pilus-specific RNA phages (Debartolomeis & Cabelli, 1991). Because of its resistance to more common tailed dsDNA phages, F^+^ HS has even been used as a marker strain to estimate water pollution via enumeration of RNA phages (Debartolomeis & Cabelli, 1991; Schaper & Jofre, 2000; Vinjé *et al*, 2004). *E. coli* HS encodes the BREX phage defense system, which has been extensively studied in heterologous context and is specific to dsDNA phages (Gordeeva *et al*, 2019; Drobiazko *et al*, 2025), suggesting encounters with tailed phages in natural environments. In addition, a native P2-like prophage DuoHS can be induced from HS chromosome and demonstrates lytic activity against prophage-free derivative (Ortiz de Ora *et al*, 2025). In summary, *E. coli* HS is widely used as a non-pathogenic control in colonization studies, yet the basis of its broad resistance to dsDNA phages, and whether this trait contributes to its success as a human gut commensal, remains unclear.

Here, we report the isolation of the first tailed phages that infect *E. coli* HS. We provide their detailed phenotypic and genomic characterization and confirm the role of the endogenous BREX system in suppressing infection. The three phage isolates were named ϕHS1, ϕHS2, and ϕHS3. They belong to the *Queuovirinae* subfamily (genus *Seuratvirus*), the *Ackermannviridae* family (genus *Taipeivirus*), and the P22-like phages (a novel candidate genus, *Hueyvirus*), respectively. Notably, ϕHS3 is a temperate phage homologous to an active endogenous HS prophage, which we named ϕHS4. Isolation of phage escape mutants revealed their tropism: ϕHS1 targets the *E. coli* HS O-antigen, while ϕHS2 and ϕHS3 target its group I capsule. Interestingly, the tail fiber proteins of ϕHS2 and ϕHS3 cluster with those of *Klebsiella* phages. Further analysis identified that the *E. coli* HS capsule was horizontally acquired from *Klebsiella* K47. Together, these results suggest that the unique combination of an O-antigen and a *Klebsiella*-like capsule, not encountered in other *E. coli* strains, makes *E. coli* HS highly resistant to coliphages, potentially contributing to its persistence in the healthy human microbiome.

## Results

### A common gut commensal *E. coli* HS is resistant to diverse tailed coliphages

To re-evaluate previously reported resistance of *E. coli* HS to tailed dsDNA coliphages (Debartolomeis & Cabelli, 1991; Schaper & Jofre, 2000), we performed an EOP assay with laboratory stock of this strain against a representative BASEL phage collection that comprises 106 phages isolated on a rough and O16-antigen restored variants of *E. coli* K12 (Humolli et al., 2025; Maffei et al., 2021). From this screen, only Bas33 produced zones of inhibited growth on HS lawns, without clearly-visible individual plaques (Fig EV1A-B). However, one-step infection growth curves revealed that Bas33 is unable to propagate on HS strain, as compared with efficient phage production on the control BW25113 host (Fig EV1C). Thus, the observed growth inhibition zones could represent the activity of viral enzymes degrading surface polysaccharides, rather than productive infection. We additionally tested sensitivity of HS to more than 70 coliphages from the laboratory collection, achieving a similar resistance profile.

To better understand the reasons for this broad phage resistance, we sequenced and annotated the laboratory variant of *E. coli* HS. The resulting assembly contains two circular components: a bacterial chromosome of 4,643,536 bp and a plasmid identical to pIMVS1_EHS (GenBank ID: CP092640.1). We compared the assembled bacterial genome to other published HS assemblies and identified several differences specific to our strain (Fig 1, Table S1), including a mutation in ribosomal protein S12 (K43T) associated with streptomycin resistance (Leatham *et al*, 2009).

**Figure 1.**
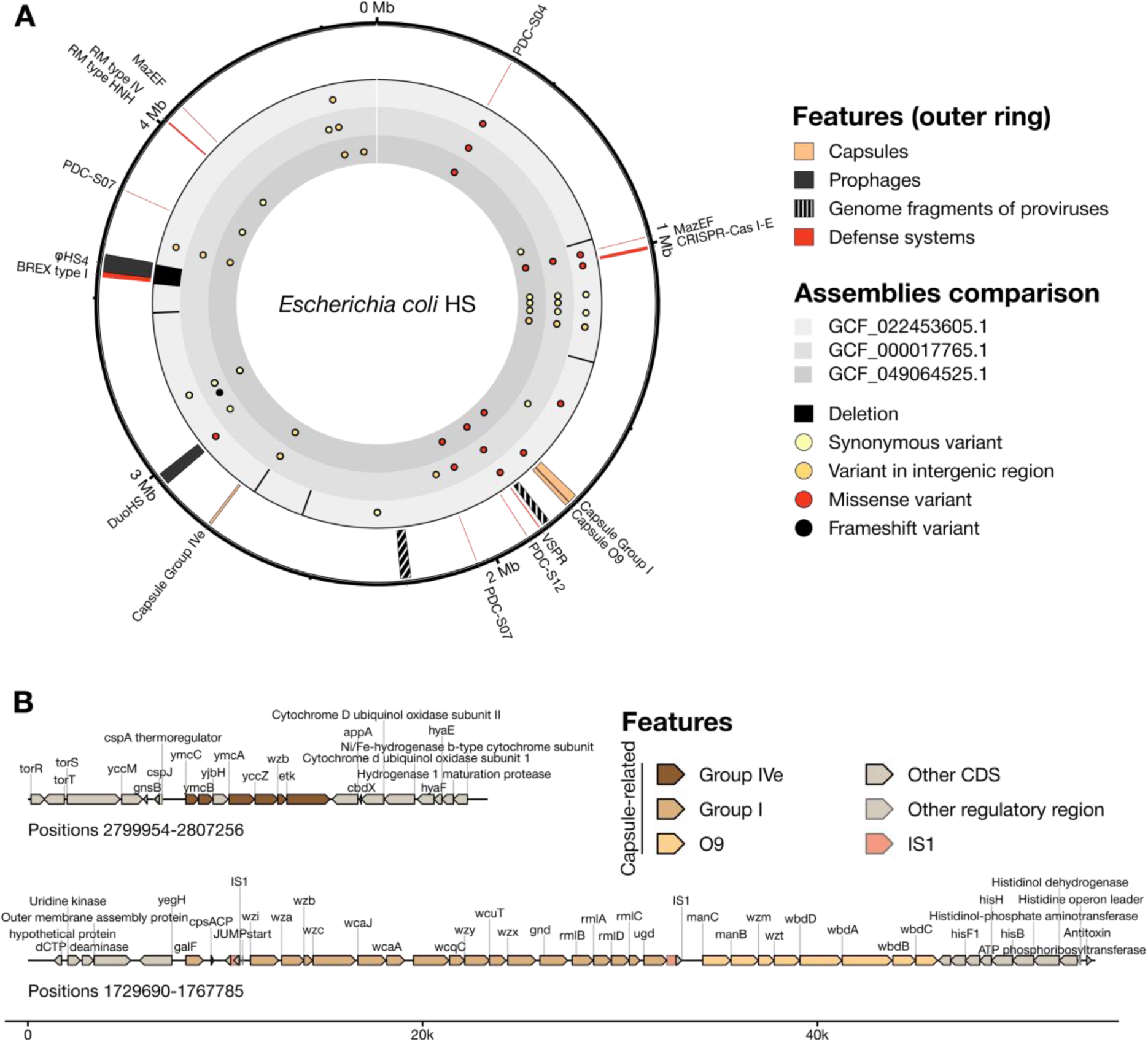
Genomic features, immune, O-antigen and capsule loci of *Escherichia coli* HS. **(A)** Circular map of the *E. coli* HS chromosome. The map highlights genetic features potentially related to phage resistance, including predicted capsule biosynthesis loci and immune system genes. Inner rings depict comparisons with publicly available assemblies of the strain (assembly accessions are provided in the legend). Single-nucleotide polymorphisms are marked with colored circles, and large-scale genomic rearrangements are indicated by filled arcs. The plasmid pIMVS1_EHS (GenBank ID: CP092640.1) is not shown. **(B)** Detailed genetic maps of the predicted capsule biosynthesis loci. The capsule loci were identified using CapsuleFinder, and gene content was manually curated based on previous studies of bacterial surface structures (Whitfield, 2006; Rasko *et al*, 2008).

Since bacterial surface structures are considered the main barrier to phage infection, we predicted capsule and O-antigen synthesis loci using CapsuleFinder (Rendueles *et al*, 2017). This analysis, followed by manual curation, revealed gene clusters related to group 4e and group 1 capsules, with the latter adjacent to the O-antigen synthesis operon (Fig 1B). Phage infection can also be inhibited by a large arsenal of immune systems or by endogenous prophages. In the HS genome, we identified two complete prophages (P2-like DuoHS and P22-like ϕHS4), two prophage remnants, and eleven immune systems predicted by PADLOC (Payne *et al*, 2022) and DefenseFinder (Tesson *et al*, 2022) (Fig 1A). Among the immune systems, only Type I BREX has been studied experimentally, and its ability to restrict infection by tailed dsDNA phages has been confirmed (Gordeeva *et al*, 2019; Drobiazko *et al*, 2025). The remaining systems are represented mostly by non-validated candidate PDC proteins.

### A novel prophage ϕHS4 is inducible but does not infect prophage-cured HS

The presence of prophages confirms previous encounters of the HS strain with tailed dsDNA phages. We therefore focused on analyzing their activity and ability to infect HS host. Among complete prophages, DuoHS belongs to the *Peduoviridae* family (Fig EV2A) and has previously been shown to be inducible (Ortiz de Ora *et al*, 2025). The second is a P22-like phage, which we denote ϕHS4. It carries a CI-like repressor with a LexA homologous domain, suggesting SOS-dependent excision (Fig EV2B) (Sauer *et al*, 1982).

To analyze whether the prophages are capable of induction and replication, we treated the HS strain with mitomycin C (MMC). Quantitative real-time PCR (qPCR) revealed a significant increase in the DNA copy number of the P22-like prophage ϕHS4, whereas the copy number of the P2-like DuoHS prophage remained unchanged (Fig EV3A). Sequencing of virion-protected DNA after MtmC induction and DNase I treatment revealed the presence of both phages in the lysate (Fig 2A). Mapping of reads to the *Escherichia coli* HS genome showed increased coverage for both prophage regions, although DuoHS coverage was approximately 100-fold lower than that of ϕHS4, consistent with the qPCR results (Fig 2A). We further attempted to obtain prophage-cured derivatives from the MtmC-treated HS culture and readily recovered a ΔϕHS4 strain (Fig EV3B), whereas multiple attempts to obtain a ΔDuoHS strain failed.

**Figure 2.**
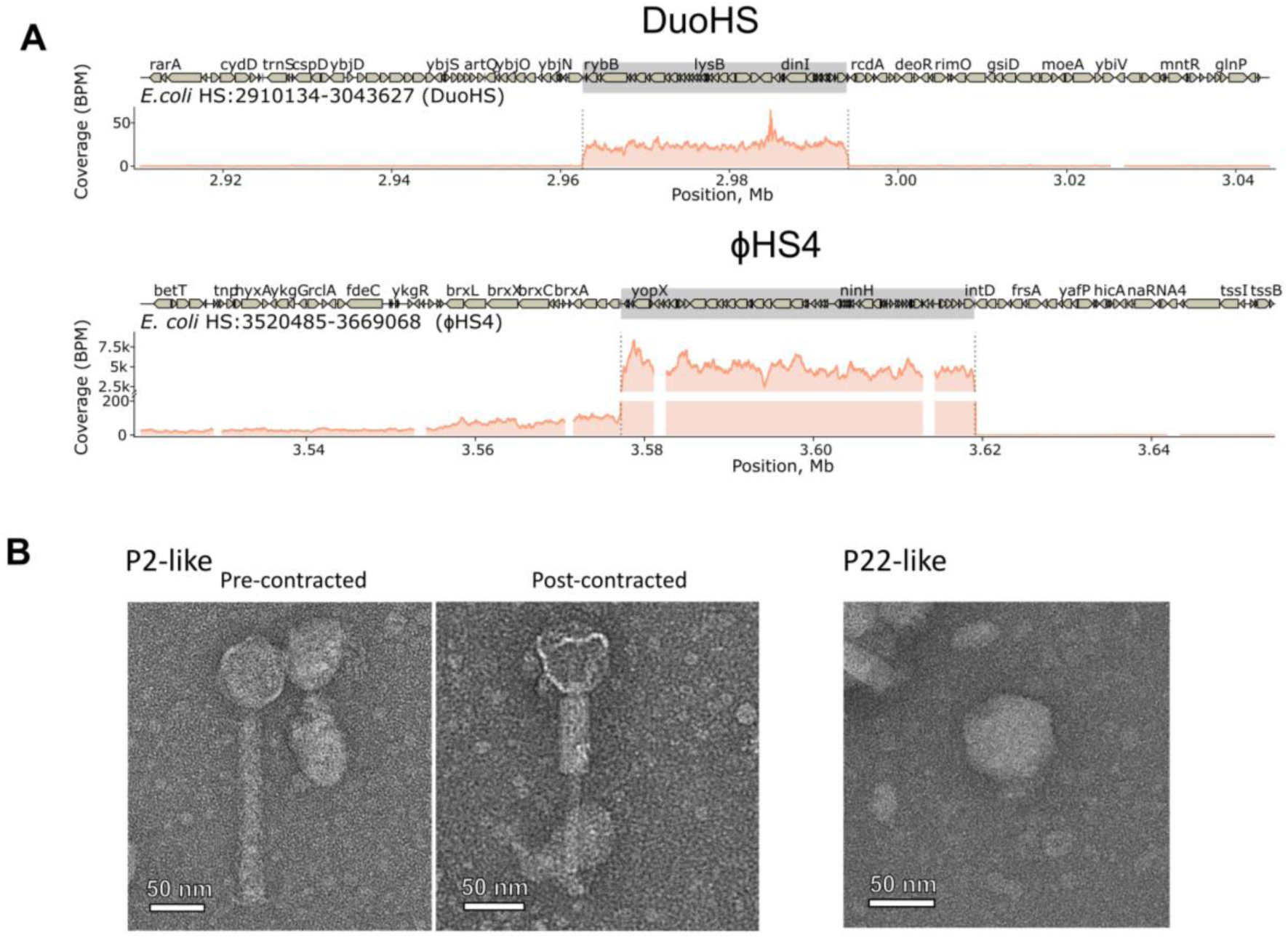
Induction and characterization of *E. coli* HS prophages. **(A)** DNA-sequencing coverage of the prophage regions of the virions that were sequenced after the induction of *E. coli* HS prophages with 0,5 μg/ml Mitomycin C (MMC). BPM (Bins Per Million mapped reads) – number of reads per bin / sum of all reads per bin (in millions). Grey regions and dotted lines indicate determined prophages. Repetitive sequences (insertion sequences) are masked for coverage, leaving gaps. **(B)** Representative TEM images of virions present in the HS lysate 3 hours after 0,5 μg/ml MMC addition. Left and centre panels show P2-like virions in pre-contracted and post-contracted states, respectively. P2-like (myovirus) morphology: an ∼86 nm head (mean 86 ± 0.6 nm) and a contractile tail of 251 ± 3.5 nm in length and 31 ± 1.2 nm in width in the extended state, shortening to ∼190 nm upon contraction. The right panel shows a P22-like virion, illustrating the characteristic morphology of this phage. P22-like (podovirus) morphology: a head diameter of ∼63.5 nm (mean 63.5 ± 1.9 nm) and a short non-contractile tail of ∼15.5 nm (mean 15.5 ± 1.4 nm). Scale bars are indicated on the images.

Transmission electron microscopy (TEM) of the MtmC-treated HS lysate revealed two distinct virion morphotypes (Fig 2B). One group exhibited P2-like (myovirus) morphology, characterized by an isometric head and a contractile tail. The second group displayed P22-like (podovirus) morphology, with a small isometric head and a short non-contractile tail. Together, NGS sequencing and TEM confirmed the ability of both prophages to form viral progeny, although the P2-like DuoHS was significantly less active in our variant of the HS strain.

To recover active phages, we plated the induced HS lysate on a range of *E. coli* hosts, including the wild-type HS strain, the ΔϕHS4 HS derivative, a prototypical K-12 strain (BW25113), and the complete ECOR reference collection (ECOR1–72) (Fig EV4A). No plaques were observed, suggesting that even the ϕHS4-cured strain remains protected against ϕHS4, either due to the presence of DuoHS or the loss of the ϕHS4 receptor following prophage acquisition. Although a zone of reduced growth was observed on the ECOR72 lawn (Fig EV4B), this was likely attributable to depolymerase activity considering the lack of productive infection. Together, these data indicate that the induced phage particles are either non-infectious or require specific host factors absent from all strains tested.

### Isolation of three novel phages that overcome the resistance of *E. coli* HS

Despite the broad resistance of the HS strain to coliphages, we hypothesized that natural reservoirs may harbor previously uncharacterized phages capable of productive infection. To bypass the active Type I BREX defense system in wild-type HS, we generated a *brxX*(Q23*) strain with a disrupted *brxX (pglX*) methyltransferase gene, thereby inactivating the BREX defense (Drobiazko *et al*, 2025). This strain was used as the sole host to screen urban waste water samples. Although the frequency of phage isolation was significantly lower than in our previous *E. coli* phage isolation campaigns, we recovered three novel bacteriophages - ϕHS1, ϕHS2, and ϕHS3 (Table 1) - and provide their detailed characterization below.

**Table 1.** Genomic features of ϕHS1, ϕHS2 and ϕHS3.

| Phage | Genome length (bp) | GC content (%) | CDS | tRNAs | (Sub)family | Genus |
| --- | --- | --- | --- | --- | --- | --- |
| ϕHS1 | 60,316 | 45% | 102 | 0 | *Queuovirinae* | *Seuratvirus* |
| ϕHS2 | 157,362 | 46% | 229 | 7 | *Ackermannviridae* | *Taipeivirus* |
| ϕHS3 | 40,197 | 47% | 71 | 0 | not classified | *Hueyvirus* |

### *Seuratvirus* ϕHS1 requires the HS O9-antigen as a receptor

Phages isolated on *E. coli* HS were plated on the ECOR collection of *E. coli* strains, which represents the natural diversity of O-antigens and receptors (Ochman & Selander, 1984; Patel *et al*, 2018). Notably, only ϕHS1 was able to infect hosts beyond HS, including the O-antigen-deficient strain BW25113 as well as ECOR 16 and ECOR 28 (Fig EV4B), encoding O9 and O104 O-antigens, suggesting that ϕHS1 has a more relaxed receptor specificity.

Transmission electron microscopy (TEM) revealed that ϕHS1 possesses a prolate head and a flexible, non-contractile tail, characteristic of siphovirus morphology (Fig 3A). On a lawn of its permissive host *E. coli* HS, ϕHS1 formed small, clear plaques (Fig 3B). A one-step growth curve at 37 °C revealed a latent period of 20–30 minutes and a burst size of approximately 100 progeny virions per infected cell (Fig 3D). Bacterial growth dynamics at low (0.001) and high (5) multiplicity of infection (MOI) were consistent with efficient lytic infection (Fig 3E). Neither the efficiency of plating (EOP), the one-step growth curve, nor the bacterial growth dynamics differed significantly between the wild-type HS (BREX+) and the HS *brxX*(Q23*) (BREX−) strains (Fig 3C–E). We conclude that ϕHS1 is not sensitive to the native BREX defense system under the conditions tested.

**Figure 3.**
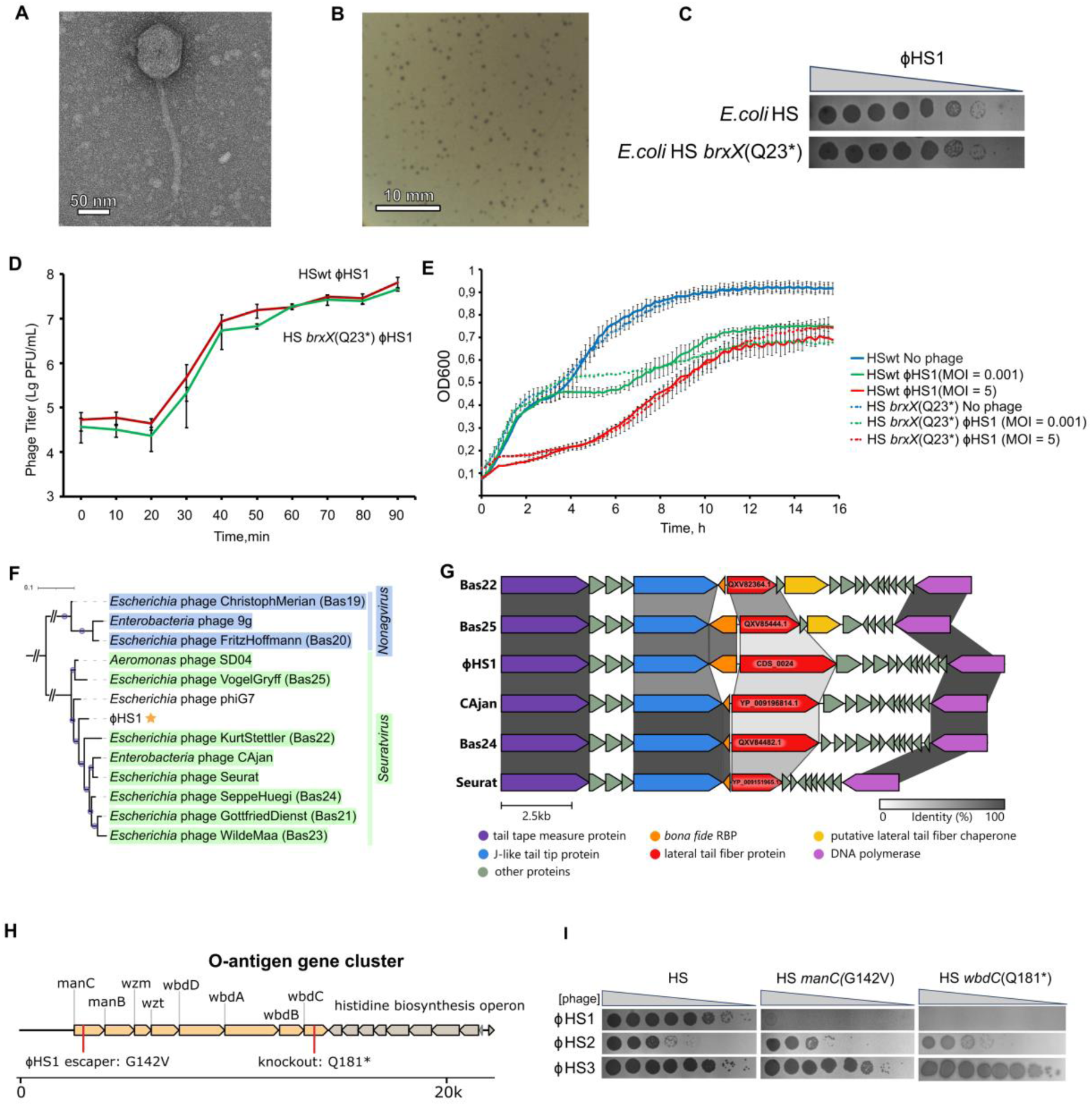
Characterization of *Seuratvirus* ϕHS1. **(A)** Representative transmission electron microscopy (TEM) image of the ϕHS1 virion. The phage has a prolate head measuring ∼79 nm in length (mean 78.5 ± 1.7 nm) × 59 nm in width (mean 58.8 ± 2.3 nm) and a flexible, non-contractile tail of ∼173 nm (mean 172.7 ± 5.6 nm). Scale bar is indicated. **(B)** Plaque morphology of ϕHS1 on an *E. coli* HS bacterial lawn. Mean plaque diameter was 0.483 ± 0.09 mm (n = 30). A representative image is shown. **(C)** Efficiency of plating (EOP) of ϕHS1 phages. EOP was assessed on lawns of wild-type *E. coli* HS and the isogenic knockout derivative HS *brxX*(Q23*), which has impaired production of the BrxX methyltransferase. **(D)** One-step growth curve of ϕHS1 in liquid LB medium at 37 °C. Infection was performed on wild-type *E. coli* HS and the isogenic HS *brxX*(Q23*) strain. Phage titers were determined at the indicated time points post-infection. Data represent mean ± SD from three independent experiments. **(E)** Growth curves of *E. coli* HS and HS *brxX*(Q23*) liquid cultures infected with ϕHS1. Cultures were infected at low (MOI 0.001) or high (MOI 5) multiplicity of infection. Uninfected controls are shown for comparison. Data represent mean ± SD from three independent experiments. **(F)** Core genome phylogeny of ϕHS1 and related *Seuratvirus* phages. The tree was constructed from protein orthogroups inferred using OrthoFinder3. The analysis included the closest related phages based on nucleotide identity, with a representative of the outgroup genus *Nonagvirus*. The tree was rooted on the outgroup branch. Branch support was assessed via bootstrapping (n = 1000); branches with support values >0.7 are indicated by blue circles. ϕHS1 is highlighted with a gold star. **(G)** Genomic alignment of the tail fiber region across *Seuratvirus* phages. The alignment of the tail fiber genomic regions of ϕHS1 and other representative *Seuratvirus* phages was generated and visualized using Clinker. Colored linkages between gene arrows represent the percentage of amino acid sequence identity of the corresponding proteins, with color intensity proportional to the identity score. **(H)** Genetic organization of the predicted *E. coli* HS O-antigen biosynthesis locus. The map shows the gene cluster responsible for O-antigen synthesis. Mutations identified in ϕHS1-resistant mutants are marked on the corresponding genes. **(I)** Efficiency of plating (EOP) of BREX-methylated ϕHS1–3 on wild-type and O-antigen mutant strains. EOP was assessed on *E. coli* HS wild-type (HS) and on a ϕHS1 resistant mutant carrying a *manC* missense mutation (G142V), and a strain carrying a premature stop codon in *wbdC* (HS *wbdC*(Q181*)), which disrupts O-antigen synthesis.

Whole-genome sequencing (Fig EV5) and phylogenetic analysis placed ϕHS1 as a novel species within the genus *Seuratvirus* (Fig 3F, Fig EV6A). The closest phage based on BLASTn total score was *Escherichia* phage phiG7, followed by *Escherichia* phage SeppeHuegi, both classified as *Seuratvirus* (subfamily *Queuovirinae*) in the NCBI taxonomy database (Turner *et al*, 2021). The intergenomic similarity between ϕHS1 and other *Seuratvirus* phages was at least 77% tANI (total Average Nucleotide Identity) (Fig EV6A), exceeding the recommended threshold of 70% for genus assignment (Turner *et al*, 2021). Phylogenetic analysis of core protein orthogroups reliably placed ϕHS1 within the *Seuratvirus* clade (Fig 3F), and major capsid protein (MCP) phylogeny fully supported this inference (Fig EV6B).

The host recognition module of ϕHS1 exhibits an organization typical of the *Queuovirinae* subfamily: it encodes a conserved tail tip protein J, a lateral tail fiber (LTF), and a non-conserved receptor-binding protein (RBP) homologous to the LptD-binding protein of phage Bas018 (Dunbar *et al*, 2025) (Fig 3G, EV6C-E).

The ϕHS1 LTF protein (CDS_0024) does not cluster with *Seuratvirus* orthologs but instead demonstrates homology to tail fibers of taxonomically diverse representatives of the genera *Tequintavirus* and *Dhillonvirus*, as well as the family *Drexlerviridae* (Fig EV6D). Notably, the ϕHS1 tail fiber is longer than those of other *Seuratvirus* tail fibers, and HHpred analysis revealed a unique C-terminal region structurally similar to the phage T5 L-shaped tail fiber protein pb1 (PDB: 4UW8) (Fig EV6F). Aligned region contains two domains: a beta-structured receptor-binding domain responsible for binding to *E. coli* O8- or O9-type O-antigen (Garcia-Doval et al, 2015), and an intramolecular chaperone, which is found in the *E. coli* phage K1F endosialidase (PDB: 3GW6, Schulz et al, 2010,Schwarzer *et al*, 2007) and the *Klebsiella* phage Kp7 tail fiber (PDB: 7Y5S), required for protein trimerization (Schulz et al, 2010). According to previous genome analysis, HS strain also encodes an O9-type antigen (Rasko et al, 2008).

The O-antigen specificity of ϕHS1 was confirmed by isolating a spontaneous ϕHS1-resistant mutant. Whole-genome sequencing revealed a G142V substitution in the *manC* gene, which encodes a bifunctional enzyme (mannose-1-phosphate guanylyltransferase and mannose-6-phosphate isomerase) required for LPS core and capsular polysaccharides biosynthesis (Fig 3H,I; Supplementary Table S2). To verify the role of the LPS O-antigen as the ϕHS1 receptor, we introduced a premature stop codon disrupting the α-1,3-mannosyltransferase gene *wbdC*(Q181*), which acts upstream of *manC* in the O-antigen biosynthetic pathway. ϕHS1 failed to infect the HS *wbdC*(Q181*) strain (Fig 3I). Notably, ϕHS1 was able to infect the O-antigen-deficient BW25113 strain (Fig EV4A), suggesting that adsorption to the O-antigen is essential only in the background of the HS strain, possibly due to reduced accessibility of the terminal receptor.

### *Taipeivirus* ϕHS2 is related to *Klebsiella*-specific phages and demonstrates tropism toward a group I capsule

The second isolated phage, ϕHS2, belongs to the genus *Taipeivirus* (family *Ackermannviridae*). TEM revealed an icosahedral head and a contractile tail bearing a branched tailspkie complex characteristic of the *Ackermannviridae* morphotype, with both pre-contracted and post-contracted states of the tail observed (Fig 4A). On a lawn of its permissive host, ϕHS2 formed small, clear plaques (Fig 4B).

**Figure 4.**
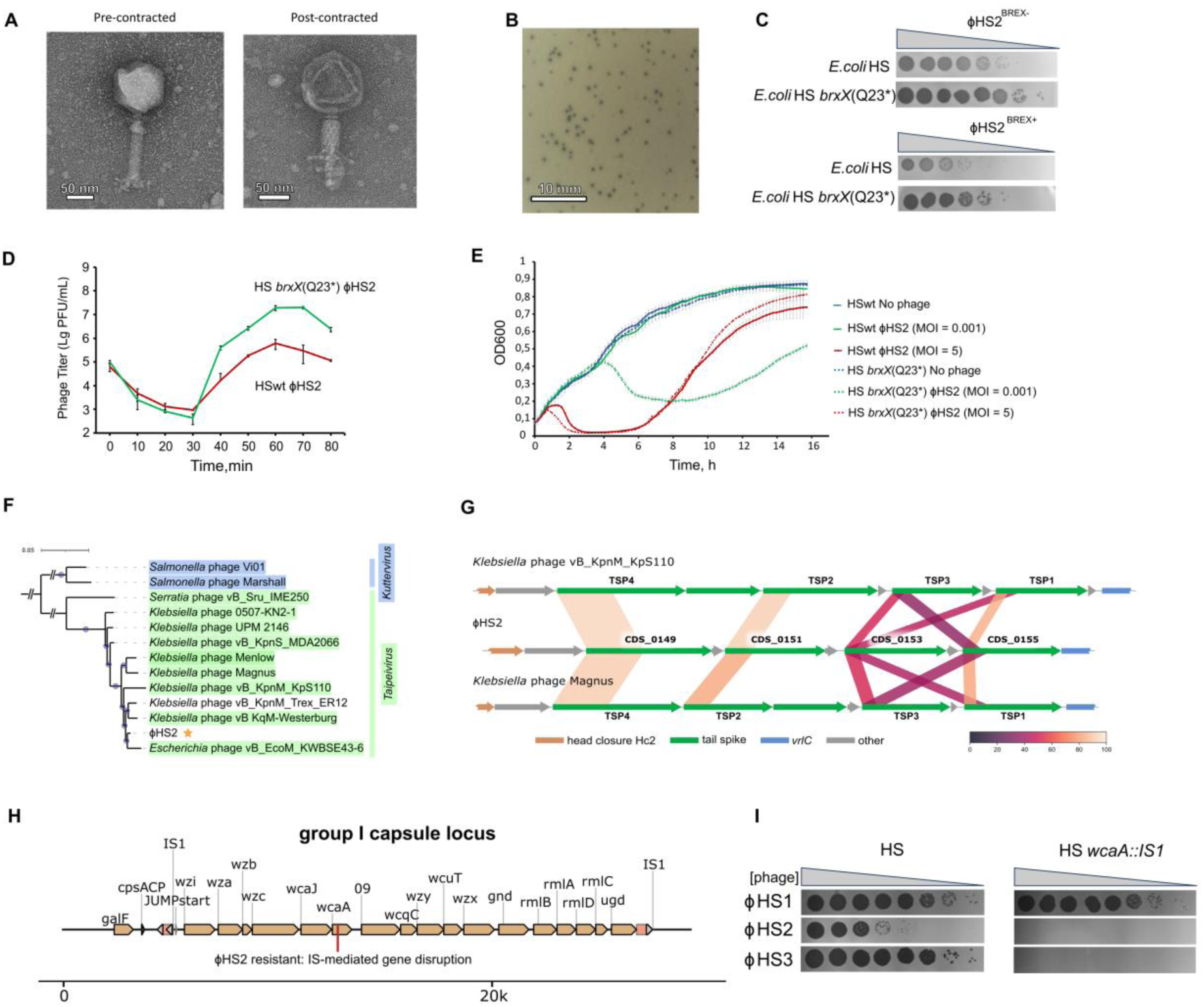
Characterization of *Taipeivirus* ϕHS2. **(A)** Representative transmission electron microscopy (TEM) image of the ϕHS2 virion. The phage displays an icosahedral head and a contractile tail characteristic of the *Ackermannviridae* morphotype. Both pre-contracted (left) and post-contracted (right) states of the tail are shown. In the extended state, the tail measured 126.8 ± 3 nm in length and 19.2 ± 0.9 nm in width; after contraction, its length decreased to 87.3 ± 16.9 nm, while its width increased to 25.6 ± 0.9 nm. Scale bars are indicated. **(B)** Plaque morphology of ϕHS2 on a lawn of the permissive host *E. coli* HS *brxX*(Q23*) (BREX⁻). Mean plaque diameter was 0.51 ± 0.10 mm (n = 30). A representative image is shown. **(C)** Efficiency of plating (EOP) of ϕHS2 on wild-type (BREX⁺) and BREX-deficient HS hosts. EOP of BREX-modified (BREX⁺) and unmodified (BREX⁻) ϕHS2 phages was assessed on lawns of wild-type *E. coli* HS (BREX⁺) and the isogenic knockout derivative HS *brxX*(Q23*) (BREX⁻), which has impaired production of the PglX methyltransferase. **(D)** One-step growth curve of ϕHS2. Infection was performed in liquid LB medium at 37 °C on wild-type *E. coli* HS and the isogenic HS *brxX*(Q23*) strain. Phage titers were determined at the indicated time points post-infection. Data represent mean ± SD from three independent experiments. **(E)** Effect of ϕHS2 infection on host cell growth. Cultures of wild-type *E. coli* HS and the HS *brxX*(Q23*) mutant were infected with ϕHS2 at low (MOI 0.001) or high (MOI 5) multiplicity of infection. Uninfected controls are shown for comparison. Bacterial growth was monitored in LB medium at 37 °C. Data represent mean ± SD from three independent experiments. **(F)** Core genome phylogeny of ϕHS2 and related phages. The tree was constructed from protein orthogroups inferred using OrthoFinder3. The analysis included the closest related phages based on nucleotide identity, with a representative of the outgroup genus *Kuttervirus*. The tree was rooted on the outgroup branch. Branch support was assessed via bootstrapping (n = 1000); branches with support values >0.7 are indicated by blue circles. ϕHS2 is highlighted with a gold star. **(G)** Genomic organization of the tail spike region across *Taipeivirus* phages. Alignment of the tail spike gene regions of ϕHS2 and other representative *Taipeiviruses*. Tail spike protein types are indicated according to a previous classification (Sørensen *et al*, 2021). Coloured linkages between the gene arrows represent the results of BLASTP local alignments, with the colour scale indicating the percentage of amino acid sequence identity. **(H)** Genetic organization of the predicted *E. coli* HS K-antigen biosynthesis locus. The map shows the gene cluster responsible for K-antigen synthesis. Mutations identified in spontaneously arising ϕHS2-resistant mutants (e.g., IS1 insertion in wcaA) are marked on the corresponding genes. **(I)** Efficiency of plating (EOP) of ϕHS2 on wild-type HS and K-antigen mutant strains. EOP of ϕHS2 was assessed on E. coli HS wild-type (HS) and on a ϕHS2 escape mutant carrying an IS1 insertion in *wcaA*, which disrupts K-antigen biosynthesis.

In contrast to ϕHS1, ϕHS2 displayed sensitivity to the BREX restriction system (Fig 4C). One-step growth curve analysis at 37 °C on the BrxX-deficient host HS *brxX*(Q23*) (BREX-) revealed a latent period of approximately 30–50 minutes and a burst size of about 100 progeny particles per infected cell (Fig 4D). On the wild-type HS (BREX+) host, however, phage burst was decreased and cell lysis at low MOI was impaired (Fig 4C–E). Of note, phage propagated on the BREX+ strain – and thus expected to acquire BREX-specific methylation – still retained BREX sensitivity, suggesting that ϕHS2 genome is non-accessible for BREX methylation (Fig 4C).

A BLASTn search of the ϕHS2 genome revealed many highly similar sequences (Fig EV7, Fig EV8). Notably, the majority of hits corresponded to *Klebsiella*-specific phages. Representatives of the genus *Taipeivirus* (family *Ackermannviridae*), including *Klebsiella* phage vB_KqM-Westerburg and *Klebsiella* phage vB_KpnM_Trex_ER12, were the closest relatives of ϕHS2 (Fig EV7, Fig EV8A). tANI values of approximately 80% are sufficient to assign ϕHS2 as a novel species within the genus *Taipeivirus* (Fig 4F, EV8). A phylogenetic tree based on protein orthogroups confirmed clustering with other *Taipeivirus* phages (Fig 4F), and a similar pattern was observed for the major capsid protein (MCP) phylogeny (Fig EV8B).

*Ackermannviridae* phages are known to possess a multicomponent tailspike protein (TSP) apparatus that determines host specificity (Sørensen & Brøndsted, 2024; Sørensen *et al*, 2024). Analysis of the ϕHS2 genome revealed a cluster of four TSP genes located between conserved *vriC* and baseplate wedge subunit genes (Fig 4G), a genomic architecture characteristic of the *Ackermannviridae* family (Sørensen *et al*, 2021). TSPs typically form homotrimers that assemble into a branched quaternary complex responsible for surface attachment and degradation of exopolysaccharide barriers (Gong *et al*, 2021; Beamud *et al*, 2023). The ϕHS2 TSPs possess a modular architecture and contain different sets of domains, with more conserved N-terminal virion attachment domains and more divergent C-terminal receptor recognition domains, also annotated as depolymerases (Fig 4G, Fig EV9).

Phylogenetic analysis revealed that ϕHS2 TSP4 (CDS_0149) is closer to the TSPs of *Klebsiella*-specific *Przondovirus* podophages than to those of other *Taipeivirus* phages (Fig EV9A,E), whereas close homologs of TSP2 (CDS_0151) were recovered from unclassified prophages of *Klebsiella* (Fig EV9B,F). The sequences of the two remaining TSPs, CDS_0153 and CDS_0155, differed significantly from *Taipeivirus* TSPs, with only short regions of local alignment, challenging their classification (Fig EV9C,D,G,H). A distinctive feature of CDS_0153 is the presence of an insert that can be annotated as a K1 capsule-specific polysaccharide lyase (IPR056204), while its C-terminal domain (PF26986) is also typical of phage depolymerases specific for *Klebsiella* capsules K3, K21, and K47 (Majkowska-Skrobek *et al*, 2018; Liu *et al*, 2020b). A BLASTP search also recovered hits for TSPs of *Przondovirus* podophages (Fig EV9C,G). Finally, CDS_0155 clustered with highly taxonomically diverse TSP homologs (Fig EV9D,H). In summary, the TSPs of ϕHS2, while showing a characteristic structural organization for *Ackermannviridae* phages, often form a separate clade with phylogenetically distinct phages and demonstrate signs of tropism toward *Klebsiella*-like capsules.

To experimentally identify the ϕHS2 receptor specificity, we isolated a spontaneous ϕHS2-resistant escape mutant of *E. coli* HS (Fig 4H,I). Short-read sequencing revealed a single persistent missense variant, R984Q, in a gene annotated as NADPH Fe³⁺ oxidoreductase subunit beta (UniRef: UniRef50_A0A6P1Q5Y9). However, we noticed that the assembly was fragmented in the group I capsular locus. Long-read (ONT) sequencing uncovered an IS1-mediated disruption of *wcaA*, a glycosyltransferase and an essential component of the capsule biosynthesis locus (Fig 4H). Notably, the resistant mutant was efficiently infected by the O-antigen-specific phage ϕHS1 but blocked plaque formation by ϕHS3 (see below), indicating that both ϕHS2 and ϕHS3 utilize the HS group I capsule as an essential receptor (Fig 4I).

### DNA hypermodifications protect ϕHS1 and ϕHS2 genomes against restriction digestion

Intrigued by the complete resistance of ϕHS1 to BREX defense and the partial resistance of ϕHS2 to both BREX defense and BREX methylation, we analyzed the ϕHS1 and ϕHS2 genomes for the presence of DNA modification loci. Phages from the *Queuovirinae* subfamily are known to substitute dG with deazaguanine derivatives, such as 2-deoxy-7-cyano-7-deazaguanine (dPreQ₀), whereas *Ackermannviridae* phages substitute dT with 5-(2-aminoethoxy)-methyluridine (5-*NeO*mdU) (Hutinet *et al*, 2021).

Similar to the previously investigated *Seuratvirus* phage CAjan, ϕHS1 encodes a conserved gene cluster responsible for the synthesis and mounting of dPreQ₀ modification (Hutinet *et al*, 2019) (Fig 5A,B). The genomic DNA of ϕHS1 was resistant or partially resistant to a set of nucleases, indicating protection at GG- and GC-containing motifs (Fig 5E), consistent with the modification pattern previously established for dPreQ₀-modified phages (Hutinet *et al*, 2019; Kot *et al*, 2020).

**Figure 5.**
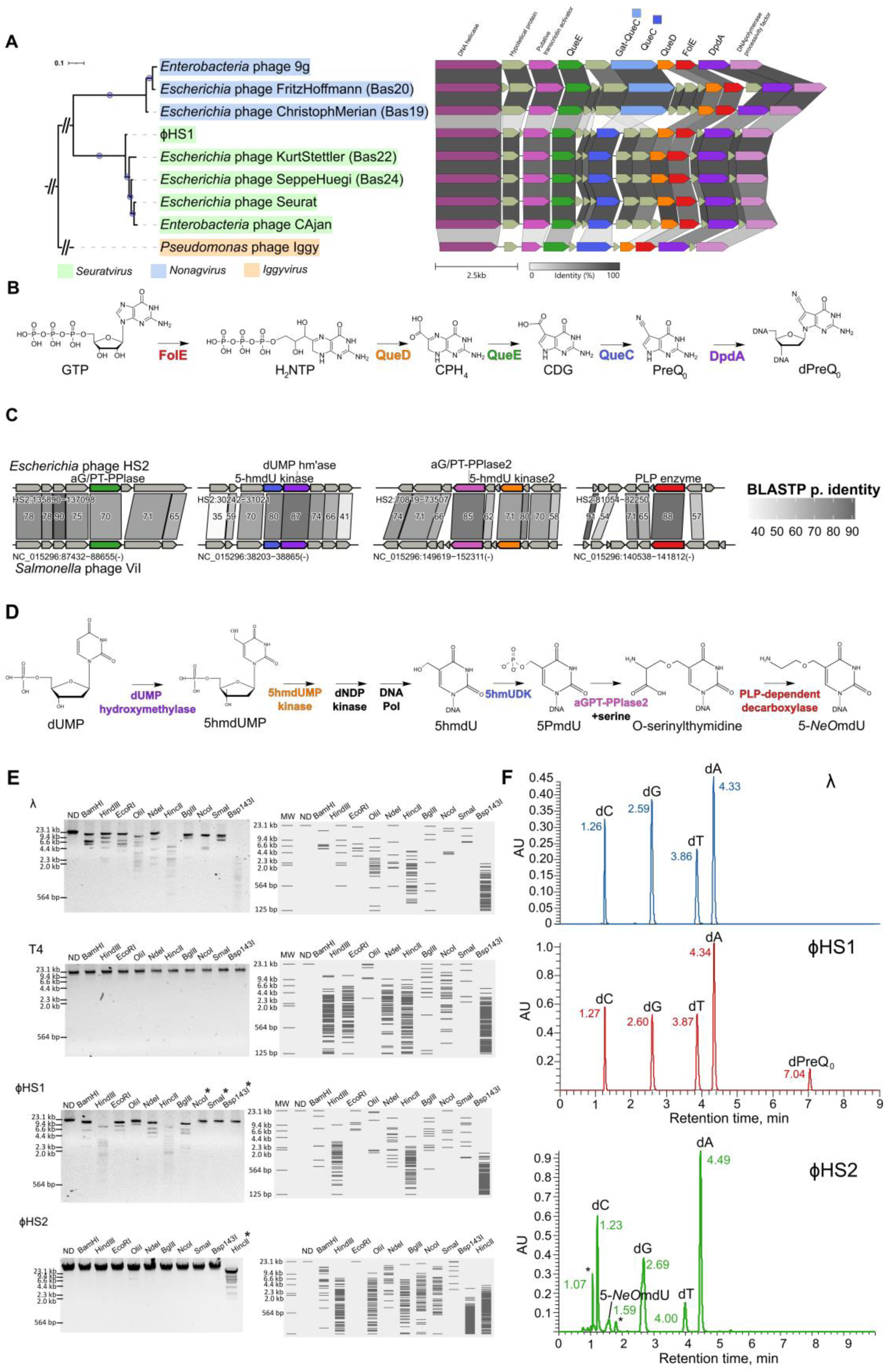
DNA hypermodifications of phages ϕHS1 and ϕHS2. **(A)** Phylogenetic placement and genomic organization of the dPreQ₀ modification cluster in ϕHS1. Left: Phylogenetic tree of ϕHS1 and selected representatives of the subfamily *Queuovirinae*, based on protein orthogroups inferred using OrthoFinder3. The tree is rooted on the outgroup (Iggy virus). Branches with bootstrap support values >0.7 are indicated by blue circles. Colored leaf backgrounds denote taxonomic groups (see legend for details). Right: Genomic alignment of the dPreQ₀ DNA modification cluster generated and visualized using Clinker. Colored linkages between gene arrows indicate the percentage of amino acid sequence identity between corresponding proteins. **(B)** Proposed biosynthetic pathway of 2′-deoxy-7-cyano-7-deazaguanosine (dPreQ₀) in the genomic DNA of ϕHS1. Abbreviations: GTP, guanosine triphosphate; H₂NTP, dihydroneopterin triphosphate; CPH₄, 6-carboxy-5,6,7,8-tetrahydropterin; CDG, 7-carboxy-7-deazaguanine; preQ₀, 7-cyano-7-deazaguanine. Enzymes: FolE (GTP cyclohydrolase I), QueD (CPH₄ synthase), QueE (CDG synthase), QueC (PreQ₀ synthase), DpdA (DNA guanine transglycosylase). **(C)** Genomic organization of the 5-*NeO*mdU modification cluster in ϕHS2 and *Salmonella* phage ViI (GenBank accession no. NC_015296.1). Genes involved in the hypermodification pathway are highlighted in bold. Colored linkages between gene arrows indicate the percentage of amino acid sequence identity between corresponding proteins. **(D)** Proposed biosynthetic pathway of 5-aminoethoxymethyl-2′-deoxyuridine (5-*NeO*mdU) in the genomic DNA of ϕHS2. Abbreviations: dUMP, 2′-deoxyuridine monophosphate; 5hmdUMP, 5-hydroxymethyl-dUMP; 5hmdU, 5-hydroxymethyl-2′-deoxyuridine; 5PmdU, 5-pyrophosphoryloxymethyl-2′-deoxyuridine. **(E)** *In vitro* restriction sensitivity assay of modified phage DNAs. Genomic DNAs from ϕHS1, ϕHS2, λ (*dam⁻ dcm⁻*) and T4 (glc-HMC) were digested with a panel of restriction enzymes. Left: Agarose gel showing experimental digestion patterns. Restriction enzymes that are evaded by the respective phage genome modifications are indicated with asterisks. Right: Expected fragment sizes (bp) for each DNA. **(F)** UV absorbance spectra (260 nm) from HPLC-MS analysis of deoxynucleosides. Spectra show the absorbance profiles of enzymatically digested genomic DNA. Labeled peaks were verified by tandem mass spectrometry (MS/MS) analysis. Asterisks indicate column artifact peaks.

Nucleosides derived from hydrolyzed ϕHS1 genomic DNA were analyzed by HPLC–MS, with bacteriophage λ gDNA serving as a non-modified control (Fig 5F). A peak unique to the ϕHS1 sample contained a compound with m/z = 292.1045, corresponding to protonated 2′-deoxy-7-cyano-7-deazaguanosine (dPreQ₀) (Fig EV10A). MS/MS analysis identified diagnostic ions at m/z = 176.1 and 159.1, matching the expected pattern of dPreQ₀ fragmentation (Fig EV10B). The level of dPreQ₀ modification was calculated from HPLC UV peak areas after correction for detector response and normalization to the phage λ signal (see Methods) (Supplementary Tables S3, S4). We estimate that 29.5% of dG residues are replaced by dPreQ₀ in ϕHS1 genomic DNA.

To determine whether dPreQ₀ modification is responsible for BREX resistance, we challenged the BREX system expressed from a plasmid in a BW25113 host against a panel of *Queuovirinae* phages (Fig EV10C). Contrary to our expectations, all phages, including ϕHS1 and 9g (which possesses a distinct deoxyarchaeosine (dG⁺) modification), were efficiently restricted by BREX (Fig EV10C). This suggests that deazaguanine derivatives do not block the recognition of GGTAAG sites by the *E. coli* HS BREX system and that ϕHS1 resistance to the endogenous BREX system of HS may be explained by insufficient expression levels of the BREX system in the native HS strain.

Inspection of the ϕHS2 genome revealed genes predicted to be involved in the biosynthesis of the hypermodified thymidine analogue 5-*NeO*mdU. Homologs of all proteins previously reported to be responsible for 5-*NeO*mdU modification in *Salmonella* phage ViI were encoded in the ϕHS2 genome (Fig 5C) (Lee *et al*, 2018). These include a dUMP hydroxymethyltransferase, 5-hmdU kinase, two aGP/TP-PPases (putative pyrophosphotransferases), and a PLP-dependent decarboxylase (Fig 5D). Accordingly, ϕHS2 genomic DNA was resistant to cleavage by all tested restriction endonucleases, with the exception of *Hinc*II (Fig 5E). We further confirmed that ϕHS2 genomic DNA contains 5-*NeO*mdU. HPLC–MS analysis of nucleosides identified a compound with the expected m/z = 302.135 (Fig 5F, Fig EV10D), and the MS/MS fragmentation pattern was consistent with 5-*NeO*mdU modification (Fig EV10E). Quantification of HPLC–MS data revealed that the modification is partial, with approximately 16.4% of dT residues replaced by 5-*NeO*mdU in ϕHS2 genomic DNA (Supplementary Table S5). Our results demonstrate that hypermodifications of ϕHS1 and ϕHS2 genomic DNA confer resistance to *in vitro* restriction digestion, whereas dPreQ₀ does not block the activity of the BREX system. Whether 5-*NeO*mdU modification interferes with the activity of BREX remains to be determined.

### ϕHS3 is a temperate P22-like phage capable of lateral transduction

TEM of the third isolated phage, ϕHS3, revealed an icosahedral head and a short non-contractile tail, characteristic of the podovirus morphotype (Fig 6A). On a lawn of the BrxX-deficient host *E. coli* HS *brxX*(Q23*), ϕHS3 formed mixed large and small plaques (Fig 6B).

**Figure 6.**
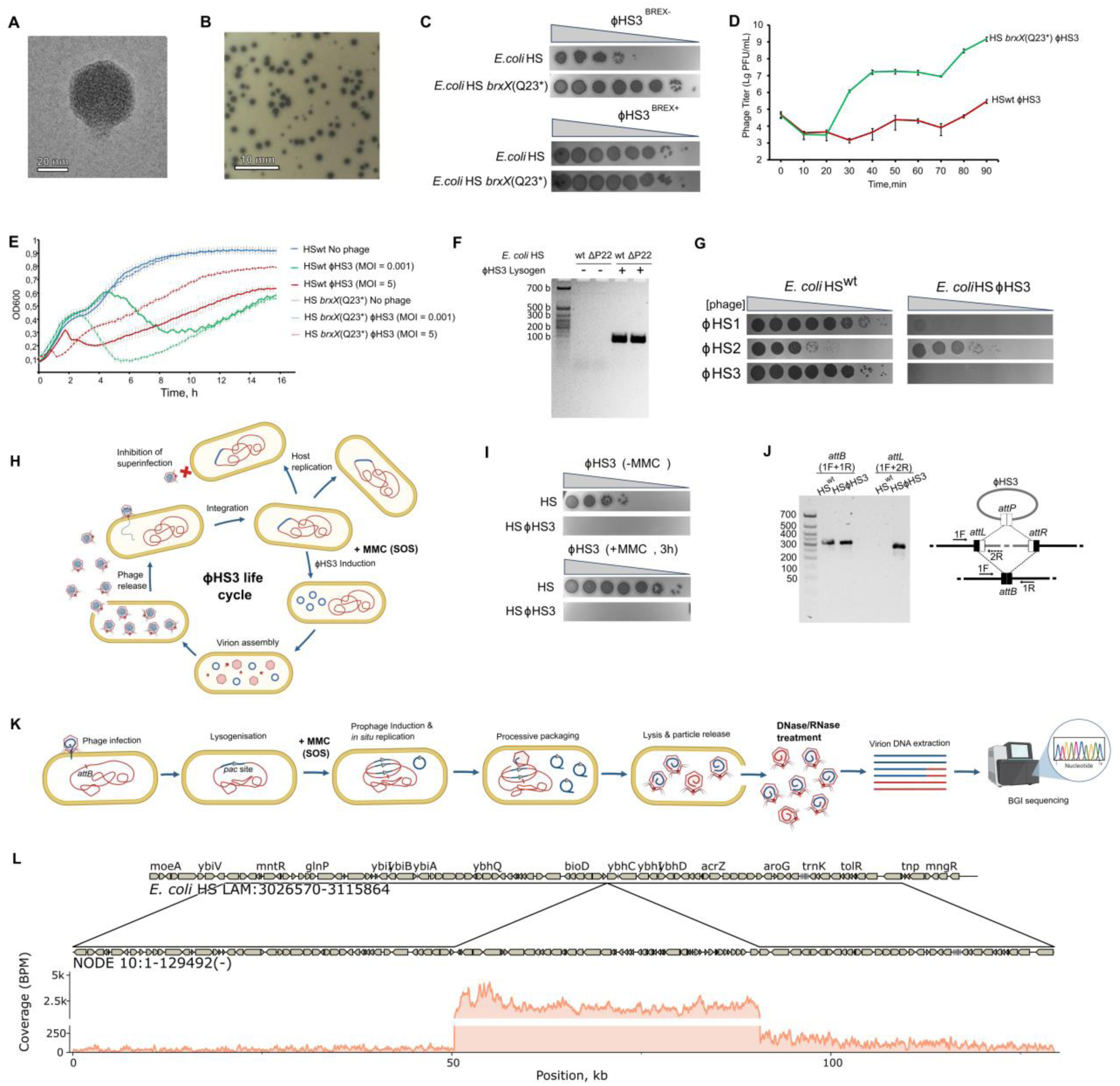
Characterization of the BREX-sensitive phage ϕHS3 and its temperate life cycle. **(A)** Representative transmission electron microscopy (TEM) image of the ϕHS3 virion. The phage displays an icosahedral head and a short non-contractile tail, characteristic of the podovirus morphotype. Head diameter is 40.8 ± 0.9 nm; tail length is 8.8 ± 0.5 nm. Scale bar is indicated. **(B)** Plaque morphology of ϕHS3 on a BREX-deficient host. Plaques of ϕHS3 on a lawn of *E. coli* HS *brxX*(Q23*) (BREX⁻). Mean plaque diameter was 1.25 ± 0.29 mm (n = 30). A representative image is shown. **(C)** Efficiency of plating (EOP) of ϕHS3 on wild-type (BREX⁺) and BREX-deficient HS hosts. EOP of BREX-modified (BREX⁺) and unmodified (BREX⁻) ϕHS3 phages was assessed on lawns of wild-type *E. coli* HS (BREX⁺) and the isogenic knockout derivative HS *brxX*(Q23*) (BREX⁻), which has impaired production of the BrxX methyltransferase. **(D)** One-step growth curve of ϕHS3. Infection was performed in liquid LB medium at 37 °C on wild-type *E. coli* HS (BREX⁺) and the isogenic HS *brxX*(Q23*) (BREX⁻) strain. Phage titers were determined at the indicated time points post-infection. Data represent mean ± SD from three independent experiments. **(E)** Effect of ϕHS3 infection on host cell growth. Cultures of wild-type *E. coli* HS (BREX⁺) and the isogenic HS *brxX*(Q23*) mutant (BREX⁻) were infected with ϕHS3 at low (MOI 0.001) or high (MOI 5) multiplicity of infection. Uninfected controls are shown for comparison. Bacterial growth was monitored in LB medium at 37 °C. Data represent mean ± SD from three independent experiments. **(F)** PCR confirmation of ϕHS3 lysogeny. PCR detection of the ϕHS3 prophage in wild-type *E. coli* HS and the HSΔϕHS4 deletion strain (lacking the P22-like prophage ϕHS4; negative controls), as well as in the newly constructed ϕHS3 lysogens HS and HSΔϕHS4. Expected amplicons are indicated. **(G)** Efficiency of plating (EOP) of BREX modified ϕHS1–3 phages on wild type and ϕHS3 lysogenic hosts. EOP was assessed on E. coli HS (BREX⁺) and the HSϕHS3 lysogenic strain. **(H)** Complete life cycle of the temperate phage ϕHS3 in *E. coli* HS. The lysogenic cycle involves prophage integration and stable maintenance. Induction of the SOS response by mitomycin C (MMC) triggers the lytic cycle, leading to prophage excision, replication, packaging and cell lysis. The experimental workflow for phage propagation and DNA isolation is shown below. (I) Efficiency of plating (EOP) of ϕHS3 lysate before and after mitomycin C induction. EOP was assessed on wild-type *E. coli* HS and a ϕHS3 lysogenic strain (HSϕHS3) using phage lysates collected either without induction (–MMC) or 3 hours after MMC treatment (+MMC). **(J)** PCR detection of the ϕHS3 genome and integration sites. Detection of the ϕHS3 prophage, as well as the bacterial attachment site (*attB*) and phage-flanking site (*attL*), in the genome of *E. coli* HS and the ϕHS3 lysogenic strain. Locus designations are indicated above the gel image. **(K)** Experimental scheme for the detection of lateral transduction. Phage lysate is induced with mitomycin C (MMC), followed by DNase/RNase treatment of released virions to remove unprotected bacterial DNA. Genomic DNA is extracted from the remaining phage particles and subjected to whole-genome sequencing. Sequencing reads are mapped back to the host bacterial genome. The schematic illustrations in panels (H) and (K) were created using <u>BioRender.com</u>. **(L)** Top: Comparison of assemblies of wild-type *E. coli* HS and the same strain harboring the ϕHS3 lysogen in the ΔϕHS4 background. Bottom: DNA sequencing coverage of the ϕHS3 integration region following induction of the ϕHS3 lysogen with mitomycin C (MtmC). BPM (bins per million mapped reads) is defined as the number of reads per bin divided by the total sum of reads per bin (in millions). Gaps in coverage indicate masked repetitive regions.

Among the isolated phages, ϕHS3 demonstrated the highest sensitivity to the BREX system in EOP assays, showing a 1,000-fold titer reduction on a BREX⁺ lawn, a phenotype that was reversed for methylated phage progeny (Fig 6C). One-step growth curve analysis on the BREX-deficient host HS *brxX*(Q23*) revealed a latent period of 20–30 minutes and a burst size of approximately 100 progeny particles per infected cell (Fig 6D). In contrast, infection of the wild-type HS (BREX⁺) host was severely impaired, as evidenced by reduced efficiency of plating and suppressed bacterial lysis (Fig 6C–E). However, eventual collapse of the BREX⁺ culture, even at low MOI, suggested a relatively rapid accumulation of methylated phages.

Genomic sequencing and annotation revealed that ϕHS3 belongs to the P22-like phages and is homologous to the endogenous *E. coli* HS prophage ϕHS4 (Fig EV11A). Based on this, we hypothesized that ϕHS3 is also capable of lysogeny. To test this, we generated a ϕHS3 lysogen in the HSΔϕHS4 background, assuming that both phages might share the same integration site (Fig 6F). However, whole-genome sequencing of the resulting ϕHS3 lysogen revealed that the integration site differed from that of ϕHS4 and was located upstream of the *ybhC* gene (Fig 6L). Moreover, ϕHS3 was also able to lysogenize the wild-type HS strain carrying the ϕHS4 prophage (Fig 6F), suggesting heteroimmunity to the ϕHS4 repressor. Nevertheless, downstream analysis of the ϕHS3 prophage was performed in the HSΔϕHS4 background.

Consistent with a temperate lifestyle, the ϕHS3 lysogen excluded ϕHS3 superinfection, as revealed by EOP assay (Fig 6G). In addition, the ϕHS3 lysogen was resistant to infection by the O-antigen-specific phage ϕHS1 but not to the capsule-specific phage ϕHS2. This phenotype could be explained by seroconversion, a phenomenon characteristic of P22-like phages. The P22 prophage expresses GtrABC proteins that modify the O-antigen structure (Allison & Verma, 2000), and ϕHS3 encodes a single protein annotated as an O-antigen ligase in the *gtrABC* locus (Fig EV11). Therefore, further research is needed to understand the mechanism of superinfection exclusion mediated by this protein.

Spontaneous excision of ϕHS3 was observed even in the absence of MtmC, as indicated by PCR with *attB*- and *attL*-specific primers and by the presence of infectious particles in the supernatant of uninduced HSϕHS3 cultures (Fig 6H-J). However, MtmC treatment resulted in a significant increase in viable phage particles, confirming that ϕHS3 exhibits a complete integration-excision life cycle and is responsive to SOS signals (Fig 6H,I). Sequencing of DNase-protected, virion-packaged DNA following MtmC induction revealed increased coverage of the corresponding ϕHS3 prophage region (Fig 6K,L). In addition, we noticed elevated coverage of the host chromosome upstream of the prophage that gradually decreased with distance (Fig 6L).

This observation is consistent with lateral transduction, a process in which packaging of the phage genome into the capsid initiates before prophage excision from the chromosome (Fillol-Salom *et al*, 2021). Since P22-like phages exploit headful packaging of concatemeric DNA, the host chromosome downstream of the *pac* site continues to be packaged into phage capsids, and this process can mobilize large fragments of the host genome. We further re-evaluated the data from MtmC induction of the wild-type HS strain and confirmed that the endogenous P22-like prophage ϕHS4 also mobilizes the adjacent chromosomal region (Fig 2A). Therefore, ϕHS3 (and possibly ϕHS4) can be used as a tool for horizontal gene transfer in *E. coli* HS.

### ϕHS3 and ϕHS4 belong to a novel phage genus, *Hueyvirus*

Phylogenetic inference for ϕHS3 proved to be more complex: its genome has low similarity to other sequences in GenBank (Fig EV12A). The maximum pairwise alignment genome coverage was 58%, albeit with high identity (∼95%). Furthermore, almost all hits were recovered from bacterial genomes, and the corresponding regions were annotated as prophages. The maximum tANI value for ϕHS3 was 55% (58% coverage) with the prophage from *Shigella flexneri* SWHIN_97, and among phage isolates, the maximum tANI was 52% (55% coverage) with *Escherichia* phage Huey (Maffei *et al*, 2025); tANI values with closest *Lederbergvirus* representatives were below 20% (Fig EV12A). The ϕHS4 genome showed even lower similarity to the closest genome: 47% tANI with the *Escherichia* KFS-D30 prophage.

These phages cannot be unambiguously classified as members of the *Lederbergvirus* genus due to the low sequence similarity. tANI below 70% is formally sufficient for the assignment of a novel phage genus. However, many phages classified by ICTV as *Lederbergvirus* also have pairwise intergenomic similarities below the genus threshold (Fig EV12A). This highlights the complexity of taxonomic assignment within P22-like phages, which is associated with their high genomic diversity and mosaicism (Casjens & Thuman-Commike, 2011).

Phylogenetic trees constructed using core protein orthogroups revealed that ϕHS3 and ϕHS4 reside in the same clade together with phages CUS-3, JEP4, vB_EcoP-720R6, and *Escherichia* prophages, whereas all ICTV-recognized exemplar isolates along with the phages Curro-Jimenez, Huey, Dewey, and Louie form a sister group to this clade (Fig.7A). Additional phylogenetic analysis was performed for major capsid proteins (MCPs), as well as for the large terminase subunit (LTS) and portal protein (PP) sequences (Fig EV12B–D). Although the topology varied slightly across protein trees, the results indicate the formation of a conserved clade comprising ϕHS3, ϕHS4, selected prophages, and phages Huey, Sf101, CUS-3, and vB_EcoP-720R6.

**Figure 7.**
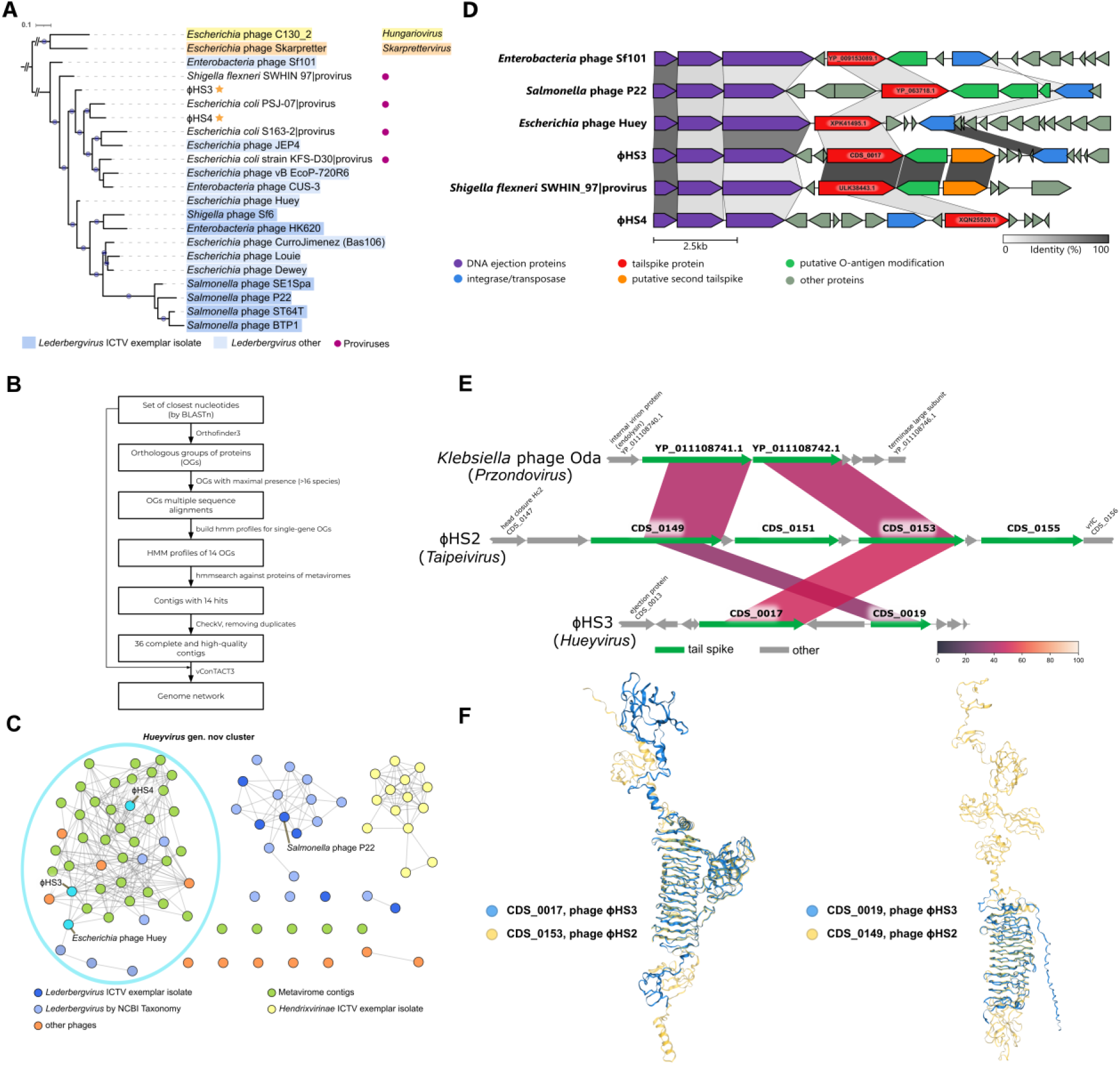
ϕHS3 belongs to a novel genus *Hueyvirus* and demonstrates a dual TSP architecture. **(A)** Core genome phylogeny of ϕHS3 and related phages within the genus *Lederbergvirus*. The tree was constructed from protein orthogroups inferred using OrthoFinder3, including the closest related genomes and representatives of the outgroup genera *Skarprettervirus* and *Hungariovirus*. The tree was rooted on the outgroup branch. Branch support was assessed by bootstrapping (n = 1000); branches with support values >0.7 are indicated by blue circles. ICTV-recognized *Lederbergvirus* exemplars are colored distinctly from NCBI-inferred taxa. Prophage regions identified by geNomad are indicated. ϕHS3 and ϕHS4 are highlighted with gold stars. **(B)** Pipeline for searching metavirome contigs based on orthologous groups of proteins from the set of closest ϕHS3-related genomes. **(C)** Genome network analysis of ϕHS3 and related metaviromic contigs. The network was constructed based on vConTACT3 analysis of a set of genomes closely related to ϕHS3 together with identified metavirome contigs. The subgraph of *Hendrixvirinae* representatives defined by vConTACT3 is shown. The network was visualized in Cytoscape (v3.10.4) after applying distance cutoff (0.51 maximum) to the edges. **(D)** Genomic alignment of tail fiber regions across *Lederbergvirus* representatives. The alignment of the tail fiber genomic regions of ϕHS3, ϕHS4 and other representative *Lederbergvirus* phages was generated and visualized using Clinker. Colored linkages between gene arrows indicate the percentage of amino acid sequence identity between corresponding proteins (linkages are not shown for “other” proteins for clarity). **(E)** Comparison of the tailspike genes regions of the ϕHS2, ϕHS3 and *Klebsiella* phage Oda (*Przondovirus* representative). Links connect regions of BLASTP local alignments for corresponding proteins; the color scale indicates the amino acid sequence identity. **(F)** Structural alignment of the ϕHS2 and ϕHS3 tailspike proteins. Shown are the AlphaFold3 models of ϕHS2 TSP (CDS_0153) and ϕHS3 TSP (CDS_0017), along with ϕHS2 TSP (CDS_0149) and ϕHS3 TSP (CDS_0019), superimposed using FoldMaso (Gilchrist *et al*, 2026).

Since the genomes available in GenBank demonstrated low similarity to ϕHS3, impeding confident taxonomic assignment, we leveraged metaviromic databases (CHDV, GDV, MGV, and HuVirDB) to extract additional genomes of ϕHS3-like phages. Using HMM profiles of previously defined orthogroups, we extracted 36 complete and high-quality contigs (Fig.7B). These genomes, along with the previously identified homologs, were compared using Vclust and taxonomically assigned using vConTACT3 (Fig 7C). The *Shigella flexneri* SWHIN_97 prophage still showed the best tANI score and the metavirome contig designated Meta_sample_29 (see Supplementary Table S6 for the original full contig names) showed a tANI of 53%.

According to the taxonomic assignments of nucleotides performed by vConTACT3, almost all phage, prophage, and metavirome-derived genomes were classified as belonging to the subfamily *Hendrixvirinae* encompassing siphoviruses, rather than podoviruses (Panigrahi *et al*, 2025). Notably, the genus *Lederbergvirus sensu stricto* has not been recognized in the current ICTV taxonomy as belonging to any taxon lower than the class *Caudoviricetes*. According to Vclust analysis and formal criteria, a total of 13 novel genera could be proposed within our dataset (Fig EV13). More importantly, vConTACT3 analysis revealed that ϕHS3, ϕHS4, Huey, and the metavirome-derived (pro)phage genomes formed a unified cluster distinct from the cluster containing classical P22 and other *Lederbergvirus* phages (Fig 7C). This result supports the designation of a novel genus, *Hueyvirus*, encompassing ϕHS3, ϕHS4, together with previously described *Escherichia* phage Huey, and retaining the same taxonomic rank as the widely recognized genus *Lederbergvirus*.

### Dual TSP architecture in ϕHS3 reveals exchange of TSPs between unrelated phages

In P22-like phages, the receptor-binding tailspike protein (gp9 in *Salmonella* phage P22 and its homologs) is encoded downstream of a conserved cluster of ejection proteins (Casjens & Thuman-Commike, 2011) (Fig 7D). In P22, gp9 recognizes repeating units of the O-antigen as a receptor but also destroys it via endoglycosidase activity (Steinbacher *et al*, 1997). TSPs of the ϕHS3 and ϕHS4 clustered with TSPs of *Klebsiella*-infecting *Przondovirus* podophages rather than with TSPs of *Lederbergvirus* phages (Fig EV14A). ϕHS3 TSP showed less than 20% similarity to the closest phage TSP, although closer homologs were recovered from *Shigella*/*Escherichia* and *Klebsiella* prophages. The set of identified ϕHS3 TSP homologs is very similar to that of TSP CDS_0153 of phage ϕHS2, and we further found that these proteins share 50% sequence identity with 81% coverage, differing only in the N-terminal virion-attachment domain (Fig 7E). This result was further supported by structural alignment (Fig 7F) and suggests that both TSPs utilize the HS capsule as a receptor.

A unique feature of ϕHS3 is the presence of a second tailspike gene (CDS_0019) (Fig 7E). A search for similar proteins recovered mostly *Klebsiella* and *Escherichia* prophage-encoded variants, followed by *Przondovirus* representatives identified previously (Fig 14B), suggesting that ϕHS3 and these phages encode related modules of two TSPs. The closest ϕHS3 homolog, however, shared only 35% identity with 80% coverage, and only 25 hits were attributed to phages in total. All of these proteins were also found among the homologs of TSP-4 of ϕHS2 (CDS_0149) and structural modeling further supports similarity of these proteins (Fig 7F). Of these homologs, TSP-3 of *Escherichia* phage Cba120 (PDB: 6NW9), was described to have glycosidase activity toward lipopolysaccharides of *E. coli* O157:H7 (Greenfield *et al*, 2019). These results suggest that ϕHS3 CDS_0019 and ϕHS2 TSP-4 may function not only as receptor-binding proteins but also as depolymerases.

Our results demonstrate unprecedented mobility of TSP-encoding genes and their exchange between P22-like phages, T7-like *Przondovirus*, and *Ackermannviridae*. Dual TSP architecture has been previously reported in T7-like podoviruses and was demonstrated to enhance their host range (Gebhart *et al*, 2017; Latka *et al*, 2019). However, to the best of our knowledge, ϕHS3 is the first P22-like phage that encodes a dual TSP host recognition complex. In addition, we reveal that *Taipeivirus* ϕHS2 and P22-like ϕHS3 encode similar capsule recognition receptor proteins that clustered with proteins from *Klebsiella*-specific phages. Consistent with this TSP similarity, an *E. coli* HS escape mutant with an IS-mediated disruption of the group I capsule locus acquired full resistance to ϕHS3 infection, confirming its capsular tropism (Fig 4H,I).

### *E. coli* HS harbors a *Klebsiella*-derived K47 capsule

Given the similarity of ϕHS2 and ϕHS3 TSPs to those of *Klebsiella*-specific phages, we re-evaluated the *E. coli* HS group I capsule structure. Kaptive annotation (Lam *et al*, 2022) revealed that the HS group I capsule closely resembles the *Klebsiella* K47 capsule, whereas the O9-antigen cluster was classified as *Klebsiella* OL3α/β. The HS K47 capsule deviated from the canonical organization owing to pseudogenization of *cpsACP* and insertion of two IS1 elements flanking the *wzi*-*ugd* cluster. Although the inverse orientation of the IS1 elements argues against their direct involvement in capsular locus mobilization, IS insertions are frequent in capsular operons and can sometimes upregulate capsule production (Wyres *et al*, 2015; Huang *et al*, 2022). Notably, an IS1 insertion at the same position as in *E. coli* HS can enhance virulence of an otherwise non-virulent *K. pneumoniae* K47 (Huang *et al*, 2022).

To test whether the HS capsular locus was acquired via horizontal gene transfer from *Klebsiella*, we searched for genetic regions similar to the HS K47/O9 locus using BLASTn against the core nucleotide database and visualized major genetic rearrangements in this region (Fig 8A). Beyond *E. coli* HS, only one strain - *Shigella flexneri* SWHIN_97, which contains a ϕHS3-related prophage (see above) - harbored a highly similar region. This suggests that *Escherichia* K47/O9 cell surface organization is extremely rare and may explain the broad resistance of HS to tailed dsDNA phages. Ten *Escherichia*/*Shigella* strains, including EC3, also encoded the K47 capsule but carried a deletion in the O-antigen synthesis locus, suggesting acquired resistance to O-antigen-specific phages. More distantly related *Escherichia* variants encoded the O9 antigen together with alternative capsules classified by Kaptive as *Klebsiella*-specific K51 or K9, pointing to multiple independent capsule exchange events between *Klebsiella* and *Escherichia*. Notably, several *Klebsiella variicola* strains exhibited a locus organization nearly identical to that of HS, including both the K47 capsule and a homologous O9/OL3α/β O-antigen synthesis cluster. The higher pairwise sequence identity between the HS and *K. variicola* K47 loci than between HS and *K. pneumoniae* suggests that *K. variicola* may have served as a source of capsule acquisition by HS strain (Fig 8A).

**Figure 8.**
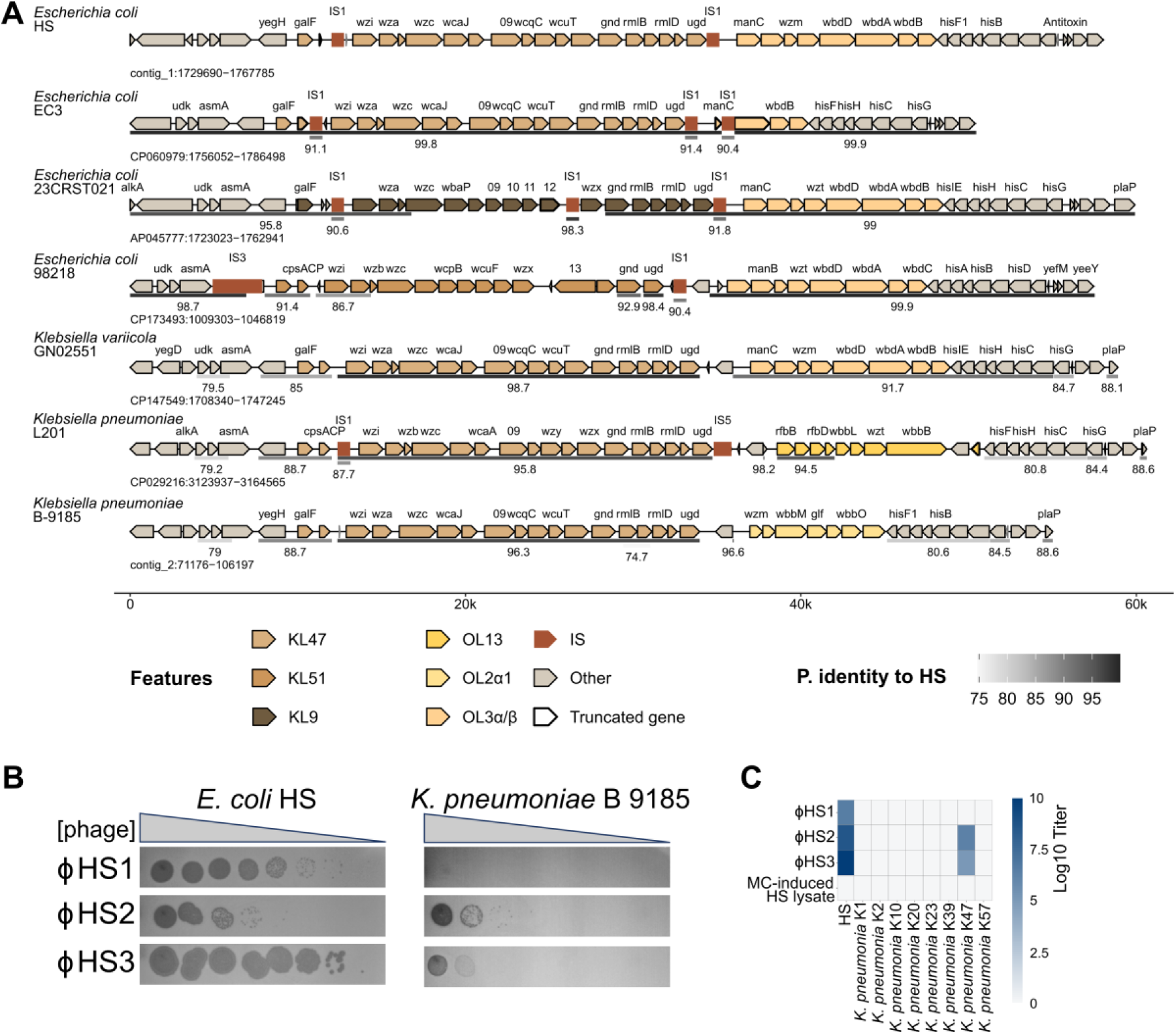
Horizontal exchange of capsular locus between *Escherichia* and *Klebsiella.* **(A)** Comparative analysis of capsule biosynthesis loci across *Escherichia* and *Klebsiella* species. Shown are the capsule loci of selected bacteria exhibiting DNA sequence similarity to the corresponding locus of *Escherichia coli* HS. The color gradient indicates the percentage of nucleotide identity of local alignments (blue to red spectrum, with red representing the highest identity). Species abbreviations: Ec – *Escherichia coli*; Kv – *Klebsiella variicola*; Kp – *Klebsiella pneumoniae*. **(B)** Efficiency of plating (EOP) of ϕHS1, ϕHS2 and ϕHS3 phages on *E. coli* HS and *Klebsiella pneumoniae* B-9185. **(C)** Heat map showing the efficiency of plating (EOP) of phages ϕHS1, ϕHS2, ϕHS3 and the mitomycin C-induced lysate from *E. coli* HS against *E. coli* HS and *Klebsiella pneumoniae* strains with different capsule types (K1, K2, K10, K20, K23, K39, K47, K57). EOP levels are color-coded as indicated.

Finally, we tested whether the K47 capsule-specific phages ϕHS2 and ϕHS3 could infect K47 *Klebsiella pneumoniae* strain B-9185. While ϕHS2 and ϕHS3 did not form plaques on any of the 72 strains from the *E. coli* ECOR collection and failed to infect *K. pneumoniae* K1, K2, K10, K20, K23, or K57 (Fig 8B,C, Fig EV4A), plaque formation was observed on the K47 strain B-9185. By contrast, the O-antigen-specific phage ϕHS1 did not infect *K. pneumoniae*. (Fig 8B). These results confirm the *Klebsiella* capsule specificity of the HS-infecting phages and demonstrate that a common *Escherichia* gut commensal, strain HS, resembles *Klebsiella* on its surface, a strategy that may help it avoid recognition by the host immune system and *E. coli*-specific gut bacteriophages.

## Discussion

The *E. coli* HS strain is often described as a true commensal of the healthy human gut and is used as a non-pathogenic control in multiple colonization studies (Leatham *et al*, 2009; Maltby *et al*, 2013; Sevrin *et al*, 2020; Winter *et al*, 2013; Elhenawy *et al*, 2019; Dalmasso *et al*, 2023; Martins *et al*, 2022; Cuenca *et al*, 2016). Remarkably, this strain has been reported to be highly resistant to dsDNA tailed phages (Debartolomeis & Cabelli, 1991; Schaper & Jofre, 2000). Our screening of 106 phages from the BASEL collection (Maffei *et al*, 2021), including phages isolated on an O-antigen-producing *E. coli* host (Humolli *et al*, 2025), confirmed this observation. Whether broad phage resistance contributes to the persistence of *E. coli* HS in the human gut remains to be determined, yet it represents a plausible hypothesis.

However, the existence of active HS prophages (P2-like DuoHS, investigated in a parallel study (Ortiz de Ora *et al*, 2025), and a P22-like ϕHS4, identified in this work) suggests that *E. coli* HS can be infected by tailed dsDNA phages in natural environments. Phage resistance can be mediated by specific cell surface structures, such as O-antigens and polysaccharide capsules, which mask terminal receptors from recognition, or by endogenous immunity proteins, which can also be encoded by resident prophages. In this work, we isolated the first tailed phages that infect *E. coli* HS and sought to determine the roles of the HS outermost surface layers (capsule and O-antigen) and the endogenous BREX immunity system in its resistance.

PADLOC identified 11 immunity-related loci in the *E. coli* HS genome (Payne *et al*, 2022). Among these, only the BREX system has been experimentally investigated previously (Gordeeva *et al*, 2019; Drobiazko *et al*, 2025). The CRISPR-Cas system is generally considered transcriptionally silent in *E. coli* (Pul *et al*, 2010), while other systems are represented by non-verified candidate PDC genes. It is likely that *E. coli* HS encodes additional, as-yet-undetermined immunity systems, including those residing in its native prophages. In this work, however, we focused on the role of the BREX system, which acts in a restriction-modification-like manner by restricting phages that lack adenine methylation at specific sites (Gordeeva *et al*, 2019; Drobiazko *et al*, 2025).

We constructed an HS derivative with a mutation in the BREX methyltransferase gene *brxX* (*pglX*), which inactivates BREX defense. Using this strain, we isolated three novel phage species: ϕHS1, ϕHS2, and ϕHS3. Analysis of the sensitivity of these phages to wild-type HS revealed that the P22-like phage ϕHS3 was severely restricted by the endogenous BREX system, while continuous phage propagation in HS culture produced methylated, BREX-resistant phage progeny. These results confirm BREX HS activity in the native background. In contrast, *Seuratvirus* ϕHS1 was completely resistant, while *Taipeivirus* ϕHS2 demonstrated only moderate sensitivity to BREX, despite carrying multiple BREX recognition sites.

Genomic analysis identified modification-related genes in ϕHS1 and ϕHS2 phages (Hutinet *et al*, 2021). Their genomic DNA was highly resistant to *in vitro* digestion by type II restriction nucleases and HPLC-MS analysis confirmed that ϕHS1 partially substitutes dG with dPreQ₀, while ϕHS2 partially substitutes dT with 5-*NeO*mdU. Strikingly, BREX expressed from a plasmid in the *E. coli* BW25113 background restricted ϕHS1 and a panel of other *Queuovirinae* phages, suggesting that dPreQ₀ modification does not limit BREX activity. The BREX system was previously reported to be inhibited by DNA mimic proteins or SAM depletion (Isaev *et al*, 2020; Andriianov *et al*, 2023). It remains possible that ϕHS1 overcomes endogenous BREX defense using a novel anti-BREX strategy. Whether 5-*NeO*mdU modification is responsible for the BREX resistance of ϕHS2 and other *Ackermannviridae* phages, similar to how glucosylation protects T4-like phages (Gordeeva *et al*, 2019), remains to be determined. In summary, although we demonstrated activity of the endogenous BREX system in *E. coli* HS, it does not represent a universal barrier to phage infection and is unlikely to be responsible for its broad phage resistance.

Further phylogenetic analysis of the receptor-binding proteins of the isolated phages revealed an interesting pattern. The tail fiber protein of ϕHS1 clustered not with *Seuratvirus* phages but rather with siphoviruses from other phage families. Lateral tail fiber proteins often enhance phage adsorption via reversible interaction with O-antigens. Isolation of an ϕHS1-resistant mutant, followed by direct inactivation of the O-antigen synthesis genetic cluster, revealed that the *E. coli* HS O-antigen is an essential receptor for ϕHS1 but does not affect ϕHS2 or ϕHS3 adsorption. Based on genomic analysis, the *E. coli* HS O-antigen has been previously serotyped as O9 (Rasko *et al*, 2008). This serotype is widespread across *E. coli* strains (Liu *et al*, 2020a) and has been reported as an essential receptor for coliphages (McCallum *et al*, 1989), interacting with their later tail fibers (Heller & Braun, 1982). Notably, it has been suggested that the O9 O-antigen can be acquired by *E. coli* from *Klebsiella*, as its structure is identical to that of *K. pneumoniae* O3 O-antigen (Vinogradov *et al*, 2002).

More intriguingly, the receptor-binding proteins of ϕHS2 and ϕHS3 have been annotated as depolymerases and cluster with proteins from phages that infect *Klebsiella*. *Ackermannviridae* phages are known to carry a branched network of tail spike proteins capable of recognizing lipopolysaccharides or capsular polysaccharides as receptors (Sørensen & Brøndsted, 2024). ϕHS2 encodes four tail spike proteins with unique C-terminal domains, all of which cluster with homologs from *Klebsiella*-specific phages. The tail spike protein of the P22-like phage ϕHS3 did not cluster with those of other P22-like phages but was surprisingly close to those of *Klebsiella*-infecting T7-like phages from the genus *Przondovirus*, revealing an extensive exchange of the TSP genes between unrelated phages. In addition, ϕHS3 seems to represent a first example of the P22-like phage with a dual TSP architecture. Notably, the TSP3 protein of ϕHS2 was similar to the tail spike protein of ϕHS3, suggesting that they share a similar tropism for the host receptor. Indeed, a ϕHS2-resistant *E. coli* HS mutant was cross-resistant to ϕHS3, but not to ϕHS1. ONT sequencing of the resistant mutant identified a group I capsular locus disruption caused by insertion of an IS1 element.

To the best of our knowledge, *E. coli* HS has not previously been described as a capsular strain, although it has been reported to have a mucoid phenotype (Schaper & Jofre, 2000) suggestive of exopolysaccharide production. Annotation of the *E. coli* HS genome with Kaptive (Lam *et al*, 2022) revealed that its group I capsule can be classified as a *Klebsiella*-specific K47 capsule. This explains the unexpected homology of ϕHS2 and ϕHS3 TSPs with *Klebsiella*-infecting phages. Furthermore, we demonstrate that ϕHS2 and ϕHS3 can cross-infect *K. pneumoniae* strain B-9185, which encodes a K47 capsule.

It has previously been noted that prophages with highly similar TSPs can be found in the genomes of *E. coli* and *K. pneumoniae* (Yang *et al*, 2024). The transfer of capsular loci from *Klebsiella* to *Escherichia* is not unique and has been reported for ∼7% of *E. coli* strains across different phylogroups (Nanayakkara *et al*, 2019). Such transfer events can even generate hypervirulent *E. coli* strains, highlighting that a change in capsular type can aid in overcoming the human immune system response (Wang *et al*, 2025). The K47 capsule of *K. pneumoniae* has also been linked to virulence (Huang *et al*, 2022), although the virulent variant contained an insertion of an IS element upstream of the capsular locus, likely increasing capsule production. It can be proposed that a change in capsular type is a highly efficient way for *E. coli* to avoid recognition by both the host immune system and bacteriophages. While the K47 capsule can be found in a few *E. coli* strains (BLASTn search identifies 12 strains with a capsular locus highly similar to HS), the combination of the *Escherichia* O9-antigen with the *Klebsiella* K47 capsule is unique for *E. coli*, and is found in a few strains of *K. variicola*, highlighting the rarity of this surface architecture, which could explain HS strain broad resistance to phages.

In addition to the three novel phages, we identified a novel active prophage in HS, which we named ϕHS4. Notably, ϕHS4 shows close homology to ϕHS3, and we further demonstrated that ϕHS3 can form lysogens and be induced in wild-type HS without interference from the ϕHS4 prophage. However, ϕHS3 lysogens became resistant to ϕHS1 infection, a phenotype that can be explained by serotype conversion of the O-antigen, mediated by the ϕHS3-encoded O-antigen ligase (Allison & Verma, 2000). P22-like phages are also known for their ability to incorporate host genomic DNA adjacent to the prophage region into phage capsids. This phenomenon, called lateral transduction, allows the mobilization of large chromosomal regions and is one of the most efficient mechanisms of horizontal gene transfer in nature (Fillol-Salom *et al*, 2021). We demonstrate that both ϕHS3 and ϕHS4 possess this capability.

The taxonomy of P22-like phages is not clearly established, owing to the mosaic nature of their genomes, which combine conserved core genes with highly variable accessory regions (Casjens & Thuman-Commike, 2011). According to formal criteria (less than 70% nucleotide identity to the closest relative), both ϕHS3 and ϕHS4 can be considered founding members of a novel genus. Given the lack of homologous phages among sequenced isolates, we leveraged metagenomic databases to identify additional (pro)phages similar to ϕHS3 and ϕHS4. vConTACT3 analysis confirmed that the metagenome-retrieved genomes, together with ϕHS3, ϕHS4, and a recently sequenced phage Huey (Maffei *et al*, 2025), constitute a separate cluster distinct from the genus *Lederbergvirus*. The genomes within the cluster still demonstrated pairwise identities below 70%, suggesting that it could be considered a (sub)family rather than a genus. However, given the current challenges in P22-like phage classification, we propose a novel group of the same taxonomic rank as the genus *Lederbergvirus*, which we name *Hueyvirus* after the first isolate from this cluster.

In summary, we isolated and characterized three novel phages infecting *E. coli* HS, a common commensal of the healthy human gut. ϕHS1 targets the O9-antigen, while ϕHS2 and ϕHS3 target the K47 *Klebsiella*-like capsule horizontally acquired from *K. pneumoniae*. This unique combination of surface structures likely explains the broad phage resistance of HS strain. Although the endogenous BREX immunity system is active in HS, it does not appear to be the major cause for this resistance. We also demonstrate that the P22-like phages ϕHS3 and ϕHS4 are capable of lateral transduction, making them potential vehicles for horizontal gene transfer. Based on their genomic distinctness, we propose the new genus *Hueyvirus* to accommodate these phages. Given the importance of phage–microbiome–host immunity interactions for human health and disease, the phages isolated in this work represent invaluable tools for future studies of commensal persistence and phage–bacteria coevolution in the gut.

## Materials and methods

### Bacterial strains, plasmids and phages

Bacterial strains and plasmids used in this study are listed in Supplementary Table S7. Bacterial cultures were routinely grown in LB medium (Luria–Bertani broth: 10 g/l NaCl, 10 g/l tryptone, 5 g/l yeast extract) at 37 °C with appropriate antibiotics where required.

*Escherichia coli* HS (laboratory stock) and its derivatives were used as host strains. The *E. coli* HS ΔϕHS4 deletion strain (lacking the P22-like prophage ϕHS4) was generated in this work. The *E. coli* HS *brxX*(*pglX*) mutant, carrying a premature stop codon in the BREX methyltransferase gene *brxX*(Q23*), and the *wbdC*(Q181*) mutant, disrupting O-antigen synthesis, were constructed using the Target-AID base editing system (Banno *et al*, 2018). Stop-knockin plasmids were assembled via BsaI-based Golden Gate assembly by ligating annealed 20-nt guide oligonucleotides into the pScI_dCas–CDA_J23119–sgRNA backbone.

Bacteriophages ϕHS1, ϕHS2 and ϕHS3 were isolated from a sample of urban waste water by direct double-layer agar plating. The water sample was cleared by centrifugation in a table-top microcentrifuge at 13 000g for 3 min and then filtered through the syringe 0.22 μm filter (Corning, Germany). 100 μL of the filtrate was plated on the lawns of *E. coli* HS and *E. coli* HS *brxX*(Q23*).

The plaques of different morphology were excised, extracted in 0.5 mL of the LB medium for 1h and serial dilutions of the extract were plated on the lawn of the corresponding isolation strain to obtain secondary plaques. The single plaque isolation procedure was repeated three times. The phages were than cultured in liquid LB medium. For growing of phage stock 3 mL of the host strain culture propagated in the shaker at 37°C and 200 rpm for 2 h (OD_600_ ∼ 0.3) and inoculated with a single plaque. The incubation was continued until visible culture lysis was observed, then 50 μL of chloroform was added. After vigorous vortexing the lysates were cleared by centrifugation as described above and the biological titer was determined.

All primers used in this study are listed in Supplementary Table S8.

### Isolation of phage and bacterial genomic DNA

Phage genomic DNA was extracted from high-titer lysates (approximately 10¹⁰ PFU/ml). Prior to DNA isolation, the lysate was treated with DNase I and RNase A (2 μl each) for 30 min at 37 °C to degrade residual bacterial nucleic acids. Phage particles were then precipitated by adding PEG 8000 (2 g) and NaCl to a final concentration of 1 M, followed by overnight incubation at 4 °C with gentle rotation. The precipitate was collected by centrifugation (30 min, ∼3,600 × g, swinging-bucket rotor) and the pellet was resuspended in 400 μl of STM buffer (50 mM Tris–HCl pH 7.5, 100 mM NaCl, 10 mM MgSO₄). Residual PEG was removed by extracting with an equal volume of chloroform. The aqueous phase was then extracted with phenol–chloroform–isoamyl alcohol (25:24:1). DNA was precipitated with 0.3 M sodium acetate and 2.5 volumes of ice-cold ethanol at −20 °C for at least 1 h, pelleted by centrifugation, washed with 70% ethanol, air-dried, and resuspended in nuclease-free water.

Bacterial genomic DNA was isolated from overnight cultures of *E. coli* HS and its derivatives using the ExtractDNA Blood & Cells kit (Evrogen, Russia) according to the manufacturer’s instructions.

### Transmission electron microscopy

For negative staining transmission electron microscopy (TEM), the formvar/carbon Cu-supported TEM grid (Ted Pella, catalog number 01801) was cleaned in Ar: O2 plasma for 40 s (1070 Nanoclean, Fischione). A volume of 20 μl of phage lysate was dropcasted onto the carbon side of the grid and left for 1 min. The residual sample was blotted by touching the grid with the blot paper, followed by two rinses in droplets of distilled H2O. After that, the grid was immediately floated on top of the drop of uranyl acetate (UA, 1 wt.% solution, 9 μl) and was held in touch with UA, droplet with tweezers for 45 s. The excess negative stain was blotted by gently sliding the side of the grid along the piece of blotting paper. The grid with the stained sample was left in the air until complete dry. Bright-field TEM images were acquired on a Titan Themis Z transmission electron microscope (Thermo Fisher Scientific) operated at 200 kV using a BM-Ceta 4 K × 4 K CMOS camera with 4-pixel binning. Virions dimensions were measured on TEM images acquired at 30,000× magnification and analyzed using ImageJ software (v1.54) (Schneider *et al*, 2012).

### One-step growth curve и EOP assays

For one-step growth curve assays, *E. coli* HS and HS *brxX*(Q23*) strains were pre-cultured in 10 ml of LB medium to an OD_600_ of 0.6 (∼2 × 10⁸ CFU/ml). Phage from a 10¹⁰ PFU/ml stock was added to achieve an initial multiplicity of infection (MOI) of 0.001. Aliquots of 1 ml were collected every 10 min, mixed with chloroform (1:100 v/v) and centrifuged at 6,000 × g for 5 min to remove bacterial cells. Phage titers in the supernatants were determined by plaque assay as described below. The experiment was performed in biological triplicate.

Phage titers were quantified using the double-layer agar overlay method. Overnight bacterial cultures (100 μl) were mixed with 10 ml of 0.6% top LB agar and poured onto pre-poured 1.2% bottom LB agar plates. Serial ten-fold dilutions of phage lysates (10 μl spots) were applied to the top agar, allowed to absorb, and plates were incubated overnight at 37 °C. For one-step growth curves, the top agar contained a 100-fold dilution of an overnight culture of *E. coli* HS *brxX*(Q23*). For efficiency of plating (EOP) assays, overnight bacterial cultures (100 μl) were mixed with 10 ml of 0.6% top LB agar with appropriate antibiotics where required. All experiments were performed in biological triplicate. Plates were imaged using an Interscience Scan1200 colony counter, and plaque diameters were measured using ImageJ (v1.54) (Schneider *et al*, 2012).

### Liquid culture infection

To monitor phage infection dynamics in liquid culture, an EnSpire Multimode Plate Reader (PerkinElmer, Hong Kong, China) was used. Overnight *E. coli* cultures were diluted 100-fold into 10 ml LB and grown at 37 °C to an OD₆₀₀ of 0.6 (∼2 × 10⁸ CFU/ml). Aliquots of 200 μl were then transferred to a 96-well plate and infected with phage from a 10¹⁰ PFU/ml stock at the indicated multiplicity of infection (MOI). Optical density was monitored continuously for 16 h. All experiments were performed in three biological replicates.

### Real-time PCR (qPCR)

qPCR assays were performed using phage-specific primers (Supplementary Table S8) in a QuantStudio 3 Real-Time PCR System (Thermo Scientific). Each 10 μl reaction contained 2 μl of 5× qPCRmix-HS (Evrogen, Russia), 500 nM primers and 1 μl of DNA (50 ng). Thermocycling conditions were: 95 °C for 5 min; 40 cycles of 95 °C for 30 s, 56 °C for 30 s, 72 °C for 30 s; followed by 72 °C for 5 min. Melt curve analysis confirmed single peaks for each primer set, indicating high specificity. Phage DNA copy numbers were quantified following 0,5 μg/ml mitomycin C induction and normalized to the host *E. coli gyrA* gene.

### HPLC-MS analysis of nucleosides

Purified genomic DNA (1–5 μg) was digested to nucleosides using Nucleoside Digestion Mix (New England Biolabs) at 37 °C overnight. Nucleosides were separated on a Waters Acquity UPLC BEH C18 column (1.7 μm, 2.1 × 100 mm) using a Waters Acquity UPLC system coupled to a Q-Exactive Orbitrap mass spectrometer (Thermo Scientific) equipped with a heated electrospray ionization (HESI) source. Separation was performed at 40 °C with a flow rate of 0.3 mL/min using 5 mM ammonium acetate (pH 4.5) as mobile phase A and 90% acetonitrile as mobile phase B. UV absorption was monitored at 260 nm. Mass spectra were acquired in positive ion mode over an *m/z* range of 200–800, and MS/MS fragmentation was performed on precursor ions of interest. Data were analyzed using Xcalibur QualBrowser (Thermo Scientific). Modified nucleosides were identified by extracted ion chromatograms, accurate mass measurements, and MS/MS fragmentation patterns (Lee & Weigele, 2021).

### Quantification of dPreQ₀ and 5-*NeO*mdU modifications

The molar amount of each canonical nucleoside was calculated from the UV peak area at 260 nm using published extinction coefficients (ε, M⁻¹·cm⁻¹): dC – 7,100, dG – 15,066, dT – 8,560, dA – 15,060 (Cavaluzzi & Borer, 2004; Palom *et al*, 2000). Since the extinction coefficients of dPreQ₀ and 5-*NeO*mdU are unknown, their quantities were determined by balance methods using the integrated peak areas (Supplementary Tables S3–S5).

For dPreQ₀ modification in ϕHS1, we assumed that the dC/dG ratio remains constant between the control phage λ and the modified phage, and that dC is unmodified. The amount of dPreQ₀ was calculated as: *n*_dPreQ₀_=*n*_dC(ϕHS1)−_*n*_dG(ϕHS1)_, where *n*_dPreQ₀_ and *n*_dG(ϕHS1)_ are the molar amounts of dC and dG in the ϕHS1 sample. The percentage of dG replaced by dPreQ₀ was then calculated by dividing the calculated amount of dPreQ₀ by the sum of dG and dPreQ₀.

For 5-NeOmdU modification in ϕHS2, we assumed that the dA/dT ratio remains constant between the control and modified phage, and that dA is unmodified. The amount of 5-*NeO*mdU was calculated as: *n*_5*NeO*mdU_=*n*_dA(ϕHS2)−_*n*_dT(ϕHS2)_, where *n*_dA(ϕHS2)_ and *n_dT(ϕHS2)_* are the molar amounts of dA and dT in the ϕHS2 sample. The percentage of dT replaced by 5-*NeO*mdU was then calculated by dividing the calculated amount of 5-*NeO*mdU by the sum of dT and 5-*NeO*mdU.

### *In vitro* digestion of bacteriophage genomic DNA

A total of 100 ng of phage genomic was incubated with indicated enzymes (BamHI, HindIII, EcoRI, OliI, NdeI, HincII, BglII, NcoI, SmaI, Bsp143I from Thermo Fisher Scientific) in commercial reaction buffer at 37 ◦C for 60 min, followed by 0.5% agarose gel electrophoresis with ethidium bromide staining.

### Oxford Nanopore Technology (ONT) sequencing

For long-read sequencing, high-molecular-weight genomic DNA was purified from overnight bacterial cultures using the Monarch Genomic DNA Purification Kit. Libraries were prepared from 500 ng of XbaI-digested DNA using the Native Barcoding Kit 24 V14 (SQK-NBD114-24) and the Ligation Sequencing Kit XL V14 (SQK-LSK114-XL) with long-fragment enrichment using the Long Fragment Buffer according to the manufacturer’s instructions. Sequencing was performed on a PromethION device using an R10.4.1 flow cell (FLO-PRO114M) with MinKNOW v23.11.2. Basecalling was performed using Dorado v0.5.3 (github.com/nanoporetech/dorado) with the super-accurate model.

### Short-read (BGI) sequencing

Libraries for whole-genome sequencing were prepared from 500 ng of phage DNA using the MGI Easy PCR-Free Library Prep Set (MGI Tech, Shenzhen, China) according to the manufacturer’s instructions. Enzymatic fragmentation was performed as recommended, followed by size selection of 400–450 bp fragments using the provided DNA Clean Beads. Library concentration was measured using a Qubit Flex fluorometer (Life Technologies, Waltham, MA, USA) with the dsDNA HS Assay Kit. Library quality was assessed with a 4200 TapeStation System (Agilent, Santa Clara, CA, USA) using the High Sensitivity D1000 ScreenTape Assay. Libraries were circularized, pooled and sequenced on a DNBSEQ-G400 platform (MGI Tech) in 2 × 150 bp paired-end mode. FastQ files were generated using ZebraCall V2 software (MGI Tech).

### *Escherichia coli* HS genome assembly

Paired short reads were trimmed and filtered via fastp v.1.01. with the following parameters: ‘--detect_adapter_for_pe --allow_gap_overlap_trimming -y -q 30’ (Chen, 2025), resulting in 14M reads. Demultiplexed long reads were filtered using Filtlong v.0.2.1 with a minimum read length of 1500, resulting in 1.5M reads with an average length of 8.3 kb (https://github.com/rrwick/Filtlong) (accessed on 1.05.2026). Hybrid assembly was performed with Unicycler v.0.5.0 with default parameters (Wick *et al*, 2017), which include short-read assembly with SPAdes v.3.15.5 (Prjibelski *et al*, 2020) (best k-mer size 127), bridging, assembly of short and long contigs with miniasm v.0.3-r179 (Li, 2016), and polishing with Racon v.1.5.0 (Vaser *et al*, 2017).

The resulting assembly contained two circular contigs, one bacterial genome (4,643,536 bp) and one plasmid (3,367 bp). This assembly was assessed for completeness using CheckM v.1.2.3 with a reduced tree (Parks *et al*, 2015). The resulting assembly of completeness 98.7, contamination 0.15 (on the marker set of *Escherichia coli*), a GC content 0.50, coverage depth >300X (determined with CoverM v0.7.0 (Aroney *et al*, 2025) and zero ambiguous bases was used for further analysis.

### *Escherichia coli* HS genome annotation

Gene prediction and primary functional annotation of *Escherichia coli* HS genome were performed with Bakta v.1.9.4 (Schwengers *et al*, 2021) with the light database (dated Feb 22, 2023). Further functional annotation was performed with default parameters of tools (unless otherwise stated). Insertion sequences were predicted with ISEScan v.1.7.2.3 (Xie & Tang, 2017). Immune systems were predicted with PADLOC v.2.0.0 (Payne *et al*, 2021) and DefenseFinder v.2.0.0 (Tesson *et al*, 2022). Capsule loci were predicted using the CapsuleFinder model set (Rendueles *et al*, 2017) with MacSyFinder v.2.1.4 (Abby *et al*, 2024) and HMMer v.3.3.2 (http://eddylab.org/software/hmmer/Userguide.pdf) (accessed on 1.05.2026). In addition, K- and O-capsules of *Klebsiella* were predicted with Kaptive v.3.1.0 (Stanton *et al*, 2025). Prophages and plasmids were demarcated with geNomad v1.11.2 (Camargo *et al*, 2024). Prophage boundaries were further revised thereafter (see Methods section: Analysis of virion DNA sequencing data after prophage induction). Antimicrobial Resistance (AMR) genes were searched with AMRFinder v.4.0.19 (database version 2024-12-18.1) for organism “Escherichia” and enabled expanded search (--plus flag) (Feldgarden *et al*, 2021).

### *Escherichia coli* HS strains genome comparison

For comparison of our in-house Escherichia coli strain HS with already deposited assemblies, search of assemblies of Escherichia coli HS in the NCBI Nucleotide database was performed (Search details: (“Escherichia coli HS”[Organism] OR Escherichia coli HS[All Fields]) OR (“Escherichia coli HS”[Organism] OR Escherichia coli strain HS[All Fields]) AND biomol_genomic[PROP]). Found genomic nucleotide sequences correspond to three assemblies (RefSeq): GCF_000017765.1 (Rasko *et al*, 2008), GCF_022453605.1, GCF_049064525.1.

The comparison of assemblies was performed by aligning them to our assembly with minimap2 v.2.26-r1175 (Li, 2018) using the parameters ‘--cs -cx asm5’. The resulting PAF files were converted to VCF with paftools.js script from minimap2 package with parameters: ‘paftools.js call -l 10 -L 100’. SNP annotation was performed with SnpEff v.5.0e (Cingolani *et al*, 2012).

### Phage genome assembly and annotation

Paired short reads were trimmed and filtered via fastp v.0.23.4 with the following parameters: ‘--detect_adapter_for_pe --allow_gap_overlap_trimming -y --dedup’ (Chen *et al*, 2018). Short-read assembly was performed with Unicycler v.0.5.0 with default parameters (Wick *et al*, 2017). All assemblies produced circular contigs, and reorientation of genomes was performed. PhageTerm v.3.0.1 (Garneau *et al*, 2017) was used but failed to produce a reoriented assembly, so homology-based reordering was utilized for the final reorientation of genomes (Korf *et al*, 2019). Completeness of phage assemblies and prophages of *Escherichia coli* HS was assessed with CheckV end_to_end pipeline (CheckV v.1.0.3 (Nayfach *et al*, 2021a)).

Gene prediction and functional annotation were performed with Sphae annotation pipeline (Sphae v.1.5.2 (Papudeshi *et al*, 2025)). Immune and anti-immune systems were predicted with DefenseFinder with parameter -a. PhageDPO (Galaxy: https://bit.ly/phagedpo, accessed on 1.11.2025, (Vieira *et al*, 2025)) was used for the prediction of depolymerase activity. For loci of modification prediction, BLASTP search was performed against a custom database of proteins involved in DNA modification (BLAST+ v.2.13.0+ (Camacho *et al*, 2009), sequences from (Kot *et al*, 2020; Lee *et al*, 2018)).

Manual curation of protein annotation was performed via search against Pfam-A database (dated of 22.01.2026) (Sonnhammer *et al*, 1997) with hmmsearch v.3.3.2 (http://eddylab.org/software/hmmer/Userguide.pdf) (accessed on 1.05.2026) and HHPRED (Söding *et al*, 2005) (access on 1.05.2026) to resolve conflicts of annotation.

### Analysis of virion DNA sequencing data after prophage induction

Short-reads DNA sequencing data after prophage induction with Mitomycin C were analysed as following. Short reads were trimmed and filtered as described above (See section *Escherichia coli* HS genome assembly). Next, reads mapping on in-house assembly was performed via BWA toolkit v.0.7.19-r1273 (Li, 2013): reference genome was indexed with ‘bwa index’, the mapping was performed with ‘bwa mem’ with default parameters. Resulting alignments were sorted and filtered with sort and view tools from SAMtools (Danecek *et al*, 2021) toolkit (filtering parameters: ‘-f 3 -q 20 -F 4,256,512,1024’). Coverage of samples sequencing was calculated with bamCoverage from DeepTools v.3.1.3 (Ramírez *et al*, 2014) package with binsize of 50 bp, and BPM normalization (BPM per bin = number of reads per bin / sum of all reads per bin in millions), excluding regions with predicted IS elements.

Based on intersection of coverage of virions DNA sequencing after prophage induction and present direct flanking repeats, boundaries of prophages were revised. To extract per-site coverage of the genome, bedtools genomecov was used (BEDtools v.2.30.0 (Quinlan & Hall, 2010)). Direct repeats were predicted with a repeat-match routine from MUMmer v.3.1 toolkit (Kurtz *et al*, 2004) with parameters specifying exhaustive search of repeats (-E) and minimal repeat length of 8 (-n 8).

To determine ϕHS3 lysogen insert site, genome of strain with deletion of ϕHS4 and lysogen of ϕHS3 was sequenced with short paired reads, and assembled with SPAdes v.4.2.0 (Prjibelski *et al*, 2020) with default parameters. ϕHS3 genome sequence was searched within the obtained fragmented assembly (N50: 83488 bp, median depth: 177.3) via minimap2 v.2.24-r1122 (Cingolani *et al*, 2012) with default parameters. To identify position of insert site within wildtype strain assembly, alignment of identified lysogen-harbouring contig on hard-masked wild-type assembly was performed. Hard-masking of reference genome was performed with genome of ϕHS3 via BBTools v.39.41 toolkit [URL: https://escholarship.org/uc/item/1h3515gn] to exclude alignment of homologous regions of phage ϕHS3 and prophage ϕHS4.

### Analysis of DNA sequencing data of phage-resistant strains

To identify genomic determinants of resistance to HS phages, both assembly and mapping of reads on the wild-type genome of *Escherichia coli* HS were performed, as described in the previous section. Variant calling was performed with the lofreq call from LoFreq v.2.1.5 (Wilm *et al*, 2012). Variants with allele frequency greater than 0.8 were annotated with SnpEff v.5.0e (Cingolani *et al*, 2012). Large-scale genomic rearrangements were searched as described in Section *Escherichia coli* HS strains comparison.

### Search for ϕHS3-like phage contigs in metavirome datasets

Four metavirome databases: Cenote Human Virome Database v.1.1 (Tisza & Buck, 2021), The Gut Virome Database v.1 (Gregory *et al*, 2020), Metagenomic Gut Virus v.1.0 (Nayfach *et al*, 2021b), and Human Virome DataBase v.1 (https://github.com/jbisanz/HuVirDB), – were downloaded and gene calling was performed with Phanotate v.1.5.1 (McNair *et al*, 2019). Protein multiple sequence alignments of 14 single-gene orthogroups specific to ϕHS3 related phages were extracted from the output of OrthoFinder. Based on these alignments, HMM profiles were built with hmmbuild from HMMer toolkit v.3.3.2 (http://eddylab.org/software/hmmer/Userguide.pdf) (accessed on 1.05.2026). Search of orthogroups within the datasets was performed with hmmsearch v.3.3.2 with E-value threshold of 1E-5. Contigs, which encode all 14 orthogroups, were extracted and filtered for completeness (completeness was assessed as described in Section Phage genome assembly and annotation). Viral contigs marked as “High-quality” and “Complete” were selected for further analysis.

### Data processing and visualization

Data processing and visualization of results was performed with Python3 v.3.11.2 [https://docs.python.org/3.11/] and R v.4.2.3 [https://www.R-project.org/] (RStudio v. 2023.3.0.386). In particular, the following packages and libraries were used: Pandas v.2.2.1, Bioframe v.0.7.2 (Open2C *et al*, 2024), NumPy v.1.26.0 (Harris *et al*, 2020), SciPy v.1.11.3 (Virtanen *et al*, 2020), Matplotlib v.3.8.0 (Hunter, 2007), Seaborn v.0.13.0 (Waskom, 2021), PyCirclize v.1.7.1 (https://github.com/moshi4/pyCirclize) (accessed on 1.05.2026), Biopython v.1.84 (Cock *et al*, 2009), Tidyverse v.2.0.0 (Wickham *et al*, 2019), gggenomes v.1.1.3 (Hackl *et al*, 2024), ggrepel v.0.9.8 (https://CRAN.R-project.org/package=ggrepel) (accessed on 1.05.2026), ggnewscale v.0.5.2.9000 [https://doi.org/10.5281/zenodo.2543762]. We aggregated the resulting figures with the help of Inkscape v.1.3.2 (091e20e, 2023-11-25) [https://inkscape.org/credits/].

### Phylogenetic and bioinformatic analysis

To identify closely related nucleotide sequences, BLASTN (v2.17.0+) online searches were performed (https://blast.ncbi.nlm.nih.gov/Blast.cgi) against the NCBI nonredundant nucleotide (nt) database (with default parameters for the program “megablast”). For the ϕHS3 phage case, more sensitive search was performed with default parameters for the program “blastn”.

Homologous protein sequences were identified using BLASTP (v2.17.0+) searches against the NCBI ClusteredNR database. Local protein sequence alignments were also performed using BLASTP.

Proviruses in bacterial genomes were predicted using geNomad v1.11.2 (‘end-to-end’ execution with parameters ‘--conservative --splits 8’) (Camargo *et al*, 2024).

Pairwise whole-genome nucleotide similarity between selected closely related genomes was calculated using Vclust v1.3.1 (parameter ‘--min-ident 0.35’ for ‘prefilter’ module, others are default) (Zielezinski *et al*, 2025). The coverage coefficients presented in heatmaps were computed as the arithmetic mean of reciprocal genome coverages, multiplied by the ratio of genome lengths.

Orthogroups were inferred using OrthoFinder v3.1.3 as well as species tree reconstruction (Emms *et al*, 2025; Emms & Kelly, 2017, 2018).

Extended taxonomic classification of phage ϕHS3, related genomes, and metavirome contigs was performed using vConTACT3 v3.1.4 (database v230) (Bolduc *et al*, 2025).

For phylogenetic analysis, amino acid sequences of major capsid proteins, large terminase subunits, portal proteins, and tail fiber/tailspike proteins were aligned using MAFFT v7.453 (Katoh & Standley, 2013). Then the maximum likelihood phylogenetic trees were constructed using the IQ-TREE v3.0.1 (Nguyen *et al*, 2015) with automatic substitution model selection by ModelFinder (Kalyaanamoorthy *et al*, 2017) and with ultrafast bootstrap option (1000 replicates) (Hoang *et al*, 2018). Trees were visualized and annotated using iTOL v7 (https://itol.embl.de) (Letunic & Bork, 2024).

Genomic regions were visualized using clinker v0.0.32 (https://cagecat.bioinformatics.nl/tools/clinker) (Gilchrist & Chooi, 2021) and pyGenomeViz v1.6.1 (https://github.com/moshi4/pyGenomeViz).

Remote protein homology detection were performed using the HHpred server (https://toolkit.tuebingen.mpg.de/tools/hhpred) (Zimmermann *et al*, 2018; Gabler *et al*, 2020).

Protein structure prediction was carried out using the AlphaFold Server (https://alphafoldserver.com) (Abramson *et al*, 2024).

Molecular graphics and analyses performed with UCSF ChimeraX v1.11.1 (Meng *et al*, 2023).

## Data availability

All materials used in this work are available upon request from the lead contact, Artem Isaev. Sequencing data generated in this study have been deposited in GenBank under accession PRJNA1467000.

## Author contributions

A. Isaev conceived the study. A. Shenfeld, O. Dorozh, O. Komarova, P. Iarema, and V. Krasilnikova performed experiments. K. Petrikov and O. Kotovskaya performed bioinformatic analysis. A. Golomidova isolated bacteriophages. A. Demkina performed sequencing. M. Zavialova performed HPLC-MS analysis. A. Isaev, N.V. Volozhantsev, K. Severinov, and A. Letarov secured resources for the study. A. Shenfeld, O. Kotovskaya, K. Petrikov, and A. Isaev wrote the manuscript. All authors reviewed and approved the final version.

## Acknowledgements

This work was supported by the Russian Science Foundation grants 24-74-00129 (to A.S.), 25-44-02137, and 24-74-10089 (to A.I.), and the Sectoral Scientific Program of Rospotrebnadzor (N.V.). The authors acknowledge the use of DeepSeek (DeepSeek, China) for grammar correction during manuscript preparation. All generated content was reviewed and approved by the authors.

