## Supplementary Data for "Phages infecting the common gut commensal *Escherichia coli* HS reveal tropism for its *Klebsiella*-like capsule"

### Appendix

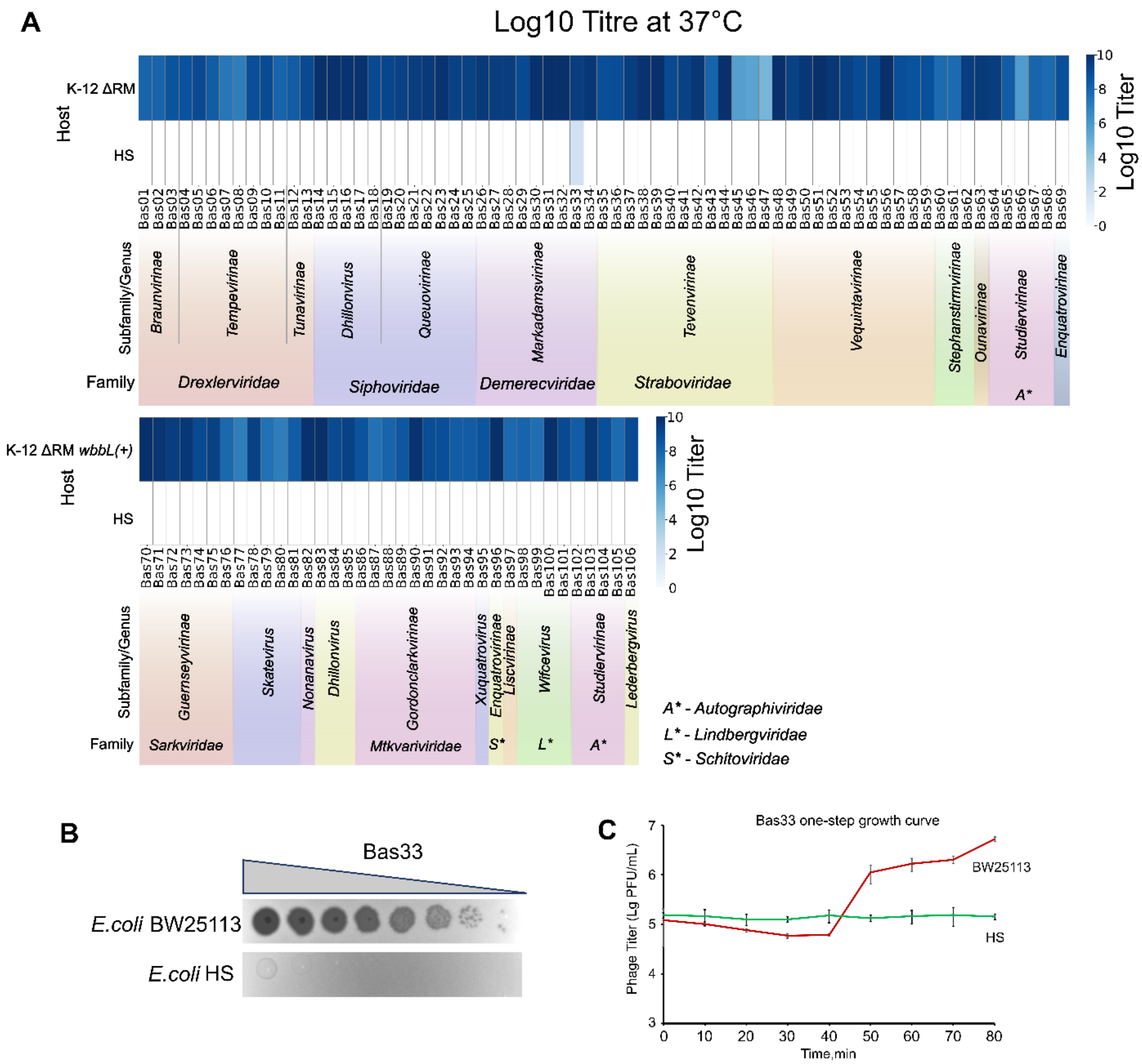


**Figure EV1. *E. coli* HS is resistant to phages from the BASEL collection. (A)** Heat map showing the infection level of *E. coli* host strains by phages Basel 1–106. For ECOR1–69, the host strain was *E. coli* K-12 ΔRM; for ECOR70–106, the host strain was *E. coli* K-12 ΔRM *wbbL*(+). **(B)** Efficiency of plating (EOP) of phage Bas33 on *E. coli* strains BW25113 (control) and HS. **(C)** One-step growth curve of bacteriophage Bas33 in liquid LB medium at 37°C. Infection was performed on wild-type *E. coli* HS and BW25113 (control).

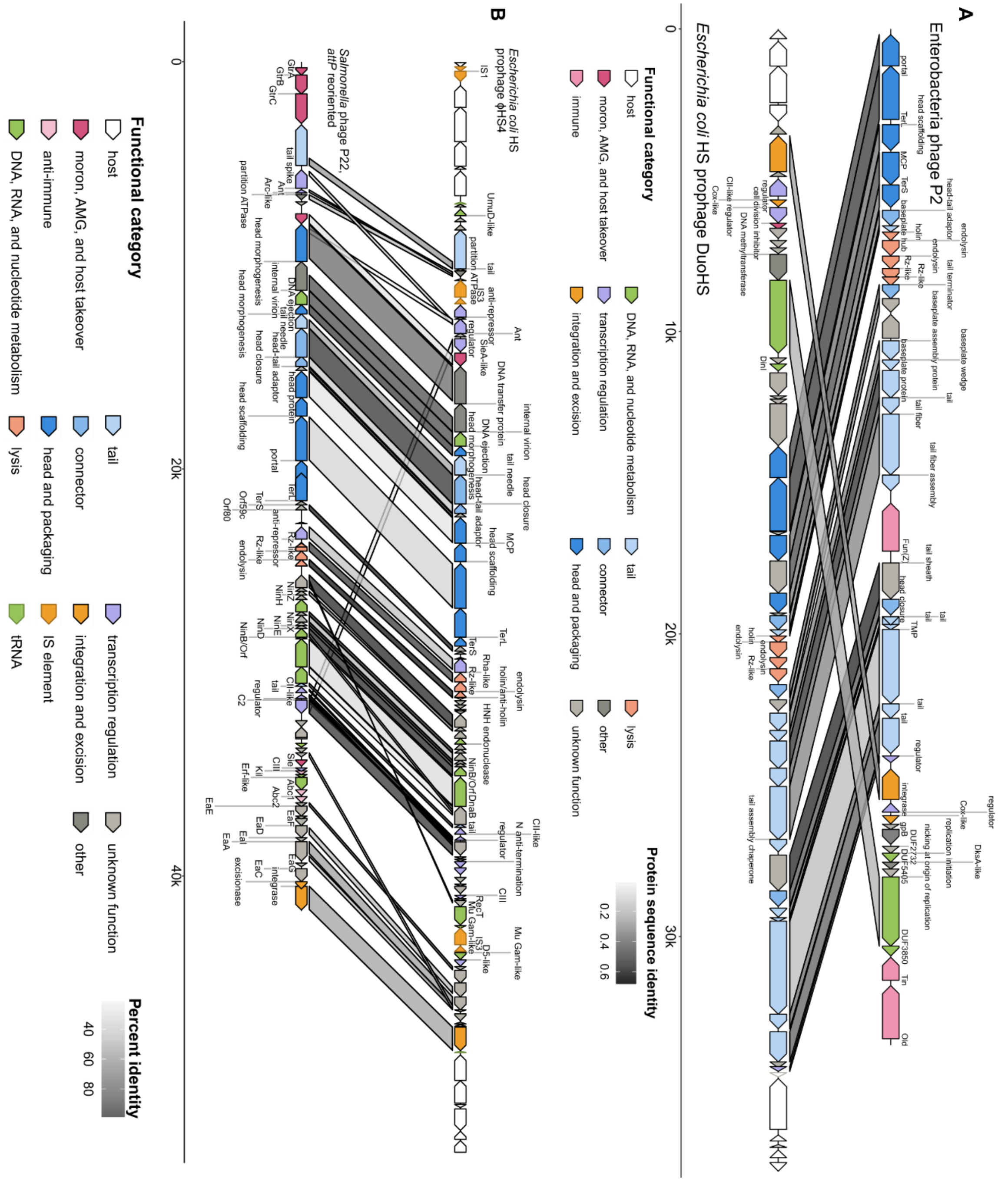


**Figure EV2. Genomic organization of *E. coli* HS prophages DuoHS and ϕHS4. (A)** Genetic map of the *E. coli* HS prophage DuoHS aligned with its reference phage P2. Genes are depicted as arrows and are color-coded according to the predicted function of their products (e.g., replication, structural, lysis). Gray shading between the maps indicates regions of protein sequence identity, with the percentage of identity shown for key genes. **(B)** Genetic map of the *E. coli* HS prophage ϕHS4 aligned with the reference phage P22 of *Salmonella* Typhimurium. Genes are depicted as arrows and are color-coded as in (A). Gray shading between the maps indicates regions of protein sequence identity, with the percentage of identity shown for key genes.

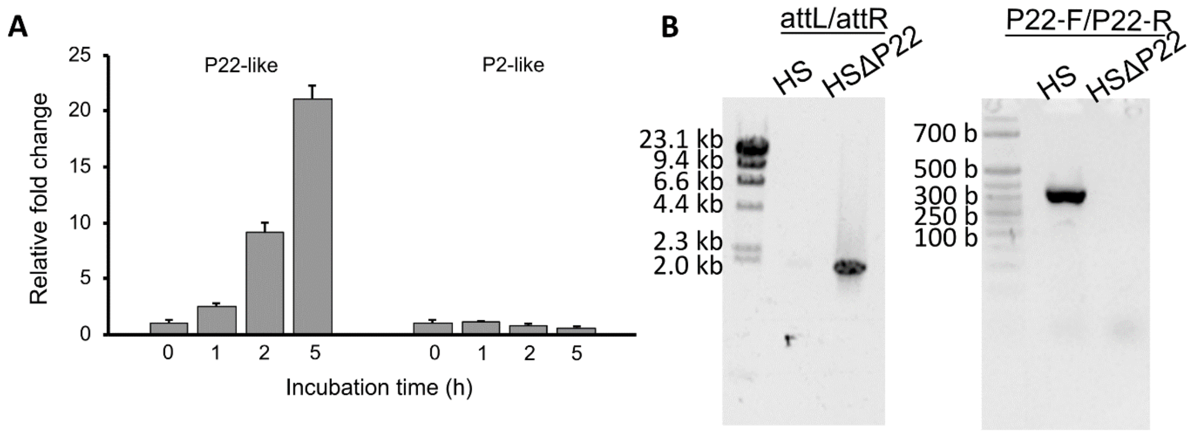


**Figure EV3. SOS-dependent induction of *E. coli* HS prophages and PCR verification of P22 prophage excision. (A)** Relative quantification of P22 (ϕHS4) and P2 (DuoHS) prophage DNA replication following SOS induction. *E. coli* HS cells were treated with mitomycin C to induce the SOS response. The change in prophage DNA copy number was determined by quantitative real-time PCR (qRT–PCR) at the indicated time points. All data were normalized to the *E. coli gyrA* gene. Data are shown as mean ± SD from three independent biological replicates. **(B)** PCR verification of P22 prophage excision (ΔP22). Left panel: PCR analysis of the phage attachment site (attB) in the wild-type HS and the ΔP22 deletion strain. In wild type HS (carrying the ~41.9 kb P22 prophage), no PCR product is detected. In the HSΔP22 strain, the expected ~1.9 kb product corresponding to the attL/attR junction is observed. Right panel: PCR analysis of the P22 prophage genome using P22-specific primers (P22-F/P22-R).

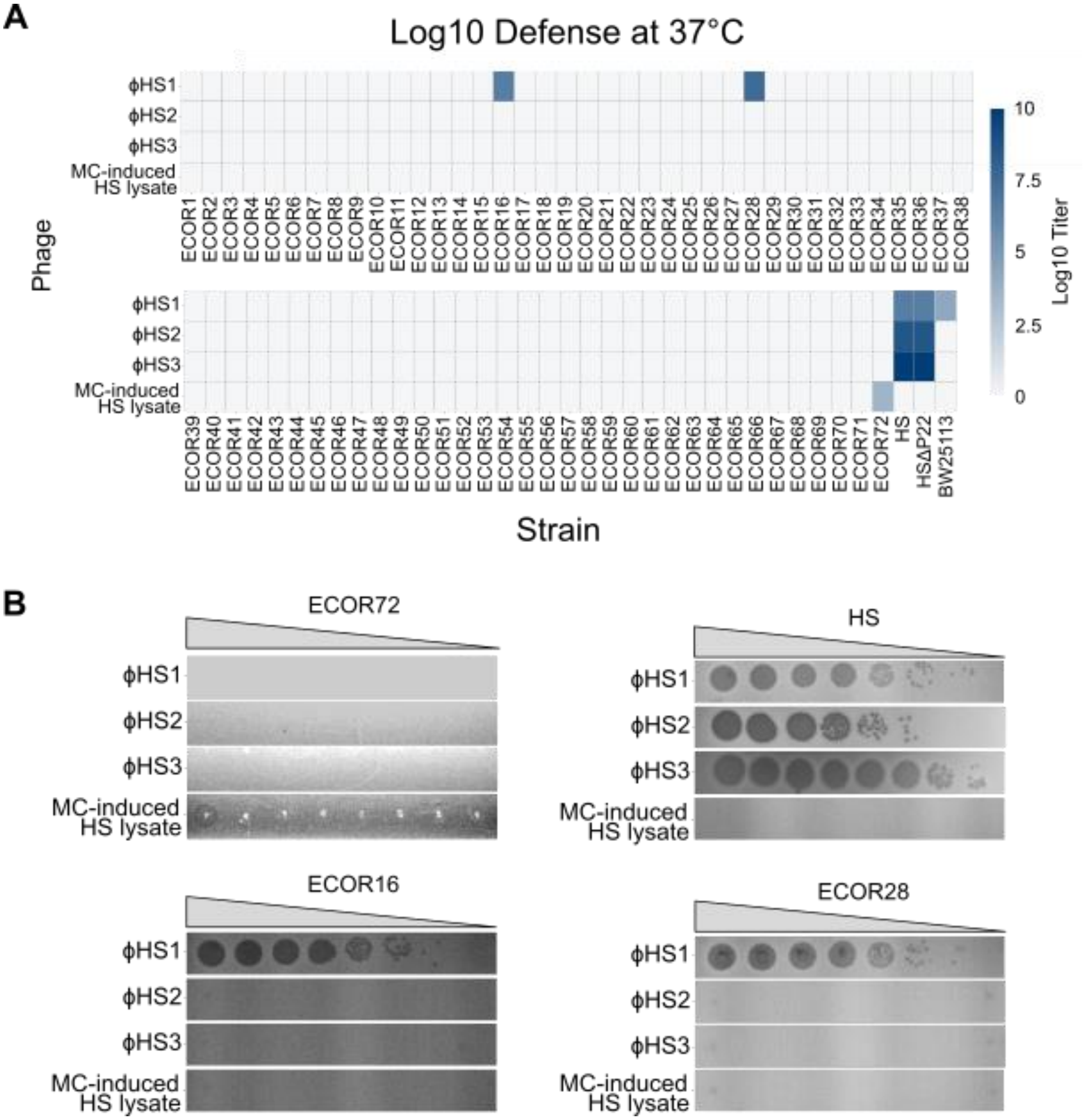


**Figure EV4. Host range of ϕHS1, ϕHS2, and ϕHS3 phages. (A)** Heat map showing the efficiency of plating (EOP) of phages ϕHS1, ϕHS2, ϕHS3 and the mitomycin C-induced lysate from *E. coli* HS against a panel of bacterial hosts: the ECOR collection (ECOR1–72), *E. coli* BW25113, *E. coli* HS, *E. coli* HSΔϕHS4. EOP levels are color-coded as indicated. **(B)** Representative images of plaque formation on selected ECOR strains. Spot assays of ϕHS1, ϕHS2, ϕHS3 and the mitomycin C-induced HS lysate on ECOR strains 16, 28, 20 and 72. A 10-fold dilution series of each phage or lysate was spotted onto bacterial lawns.

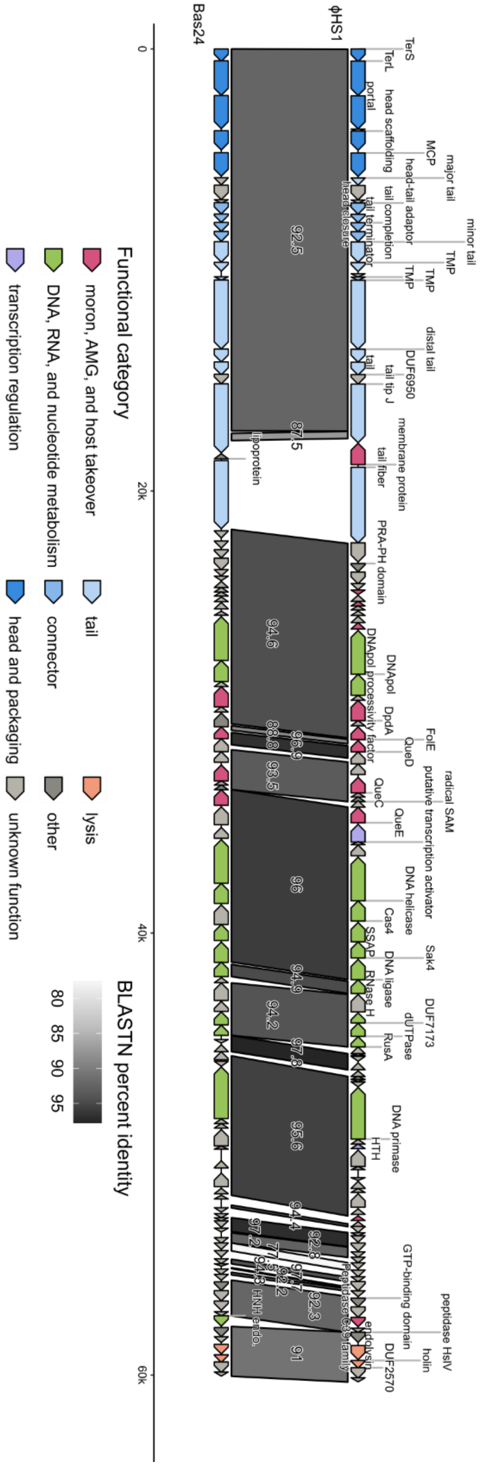


**Figure EV5. Genomic organization of *Seuratvirus* phage ϕHS1.** Comparative genome organization of ϕHS1 and the related *Seuratvirus* phage SeppeHuegi (Bas24). Arrows represent genes and are color-coded according to predicted function. Gray shading between the genomes indicates regions of DNA sequence similarity.

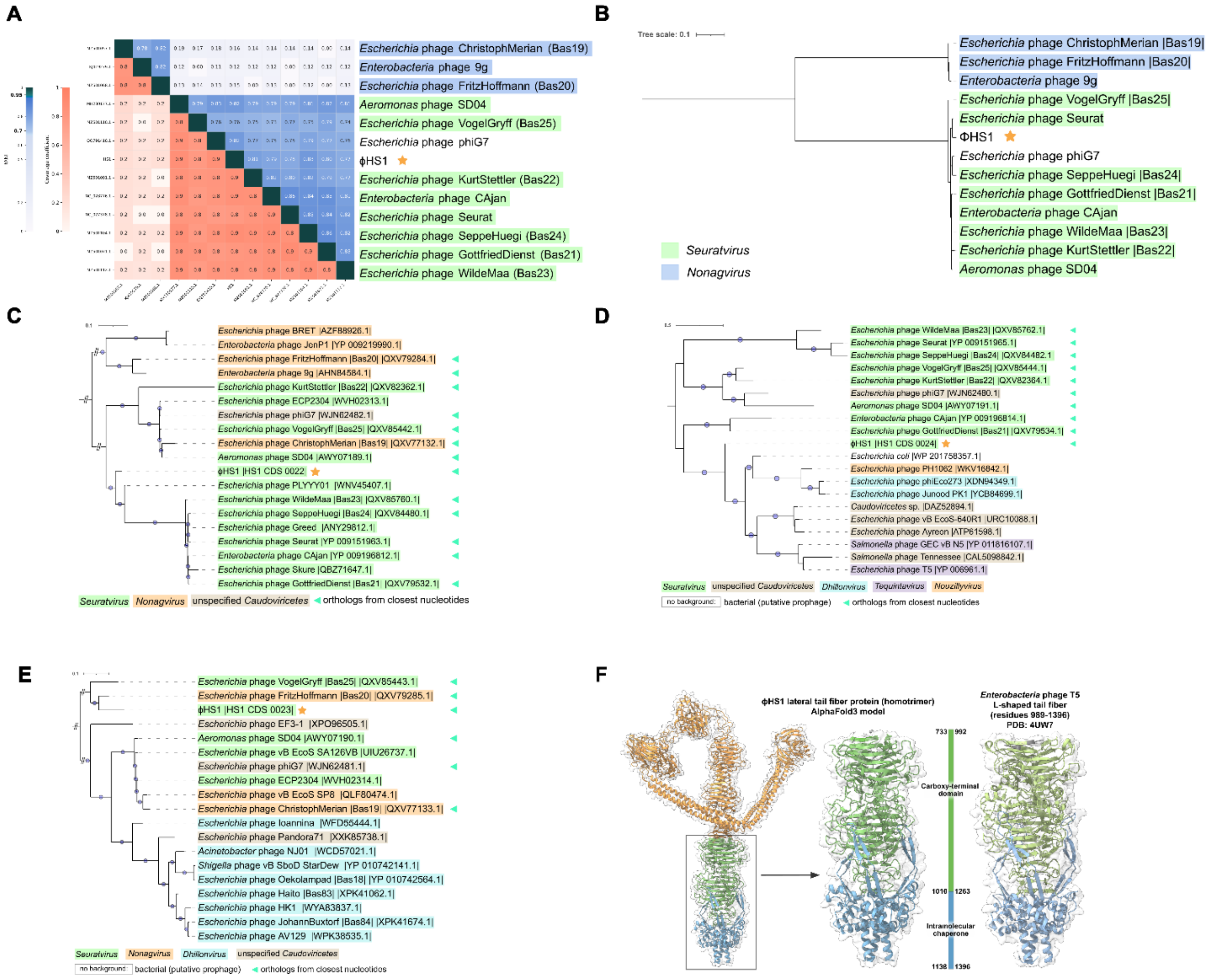


**Figure EV6. Nucleotide identity and phylogenetic analysis of ϕHS1 and its receptor binding proteins. (A)** Whole-genome nucleotide identity matrix of ϕHS1 and related phages. The matrix shows the average nucleotide identity (ANI) between each pair of genomes. A representative of the outgroup genus *Nonagvirus* is included. **(B–E)** Midpoint-rooted maximum likelihood phylogenetic trees of key ϕHS1 proteins. Trees are shown for: **(B)** major capsid protein (MCP); **(C)** J-like tail tip protein (CDS_0022); **(D)** lateral tail fiber (CDS_0024); and **(E)** *bona fide* receptor-binding protein (RBP; CDS_0023). Colored backgrounds indicate taxonomic groups, with bacterial labels representing NCBI Identical Protein Groups. ϕHS1 is highlighted in each tree with a gold star. **(F)** Structural comparison of the ϕHS1 lateral tail fiber protein with the T5 phage tail fiber. The left panel shows the AlphaFold3 model of the ϕHS1 lateral tail fiber homotrimer (entire complex on the left; enlarged C terminal part in the center). The right panel shows the structure of the Enterobacteria phage T5 tail fiber (PDB: 4UW7). Domains are colored according to the color bar, with numbers indicating the boundaries of the regions aligned by HHpred, shown as amino acid positions in the corresponding proteins.

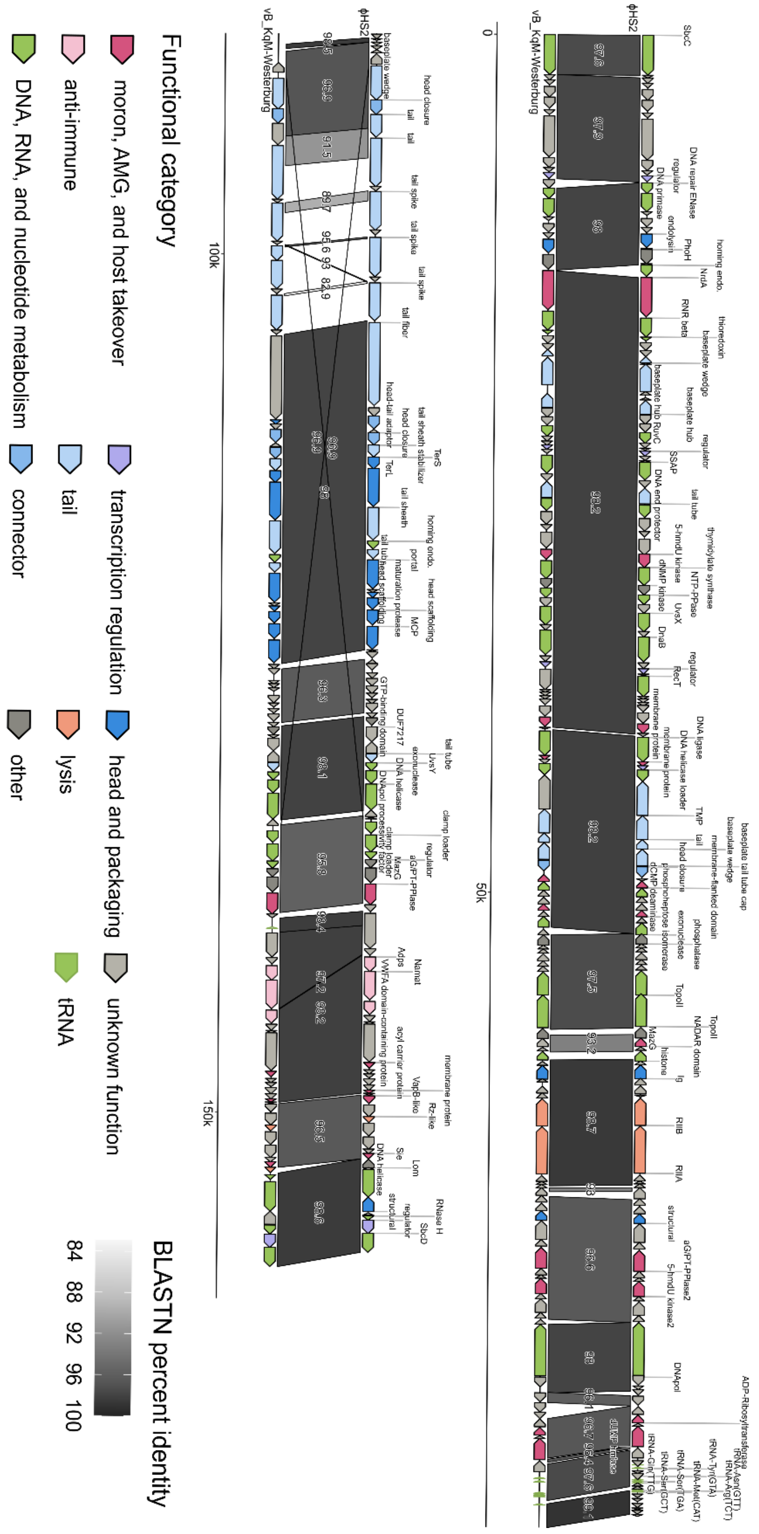


**Figure EV7. Genomic organization of *Taipeivirus* phage ϕHS2.** Comparative genome organization of ϕHS2 and the related *Taipeivirus* phage vB_KqM-Westerburg. Arrows represent genes and are color-coded according to predicted function. Gray shading between the genomes indicates regions of DNA sequence similarity.

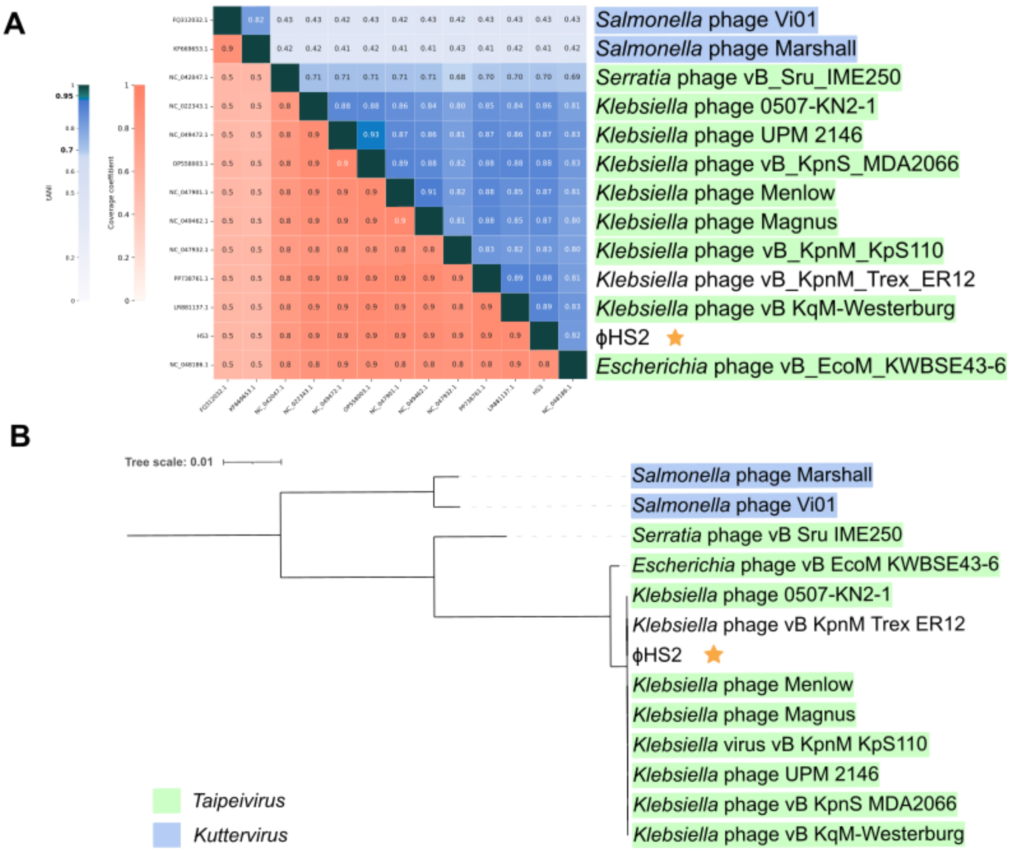


**Figure EV8. Taxonomic classification of ϕHS2. (A)** Whole-genome nucleotide identity matrix of ϕHS2 and related phages. The total average nucleotide identity (ANI) matrix was constructed using Vclust for the set of closest related genomes, with representatives of the outgroup genus *Nonagvirus* included. **(B)** Phylogenetic tree of the ϕHS2 major capsid protein (MCP). A midpoint-rooted maximum likelihood tree is shown. Homologs were obtained from the closest nucleotide orthogroup (green triangles) and by BLASTP searches. Background color indicates the lowest taxonomic rank; bacterial protein labels denote the organism group corresponding to the NCBI Identical Protein Groups entry.

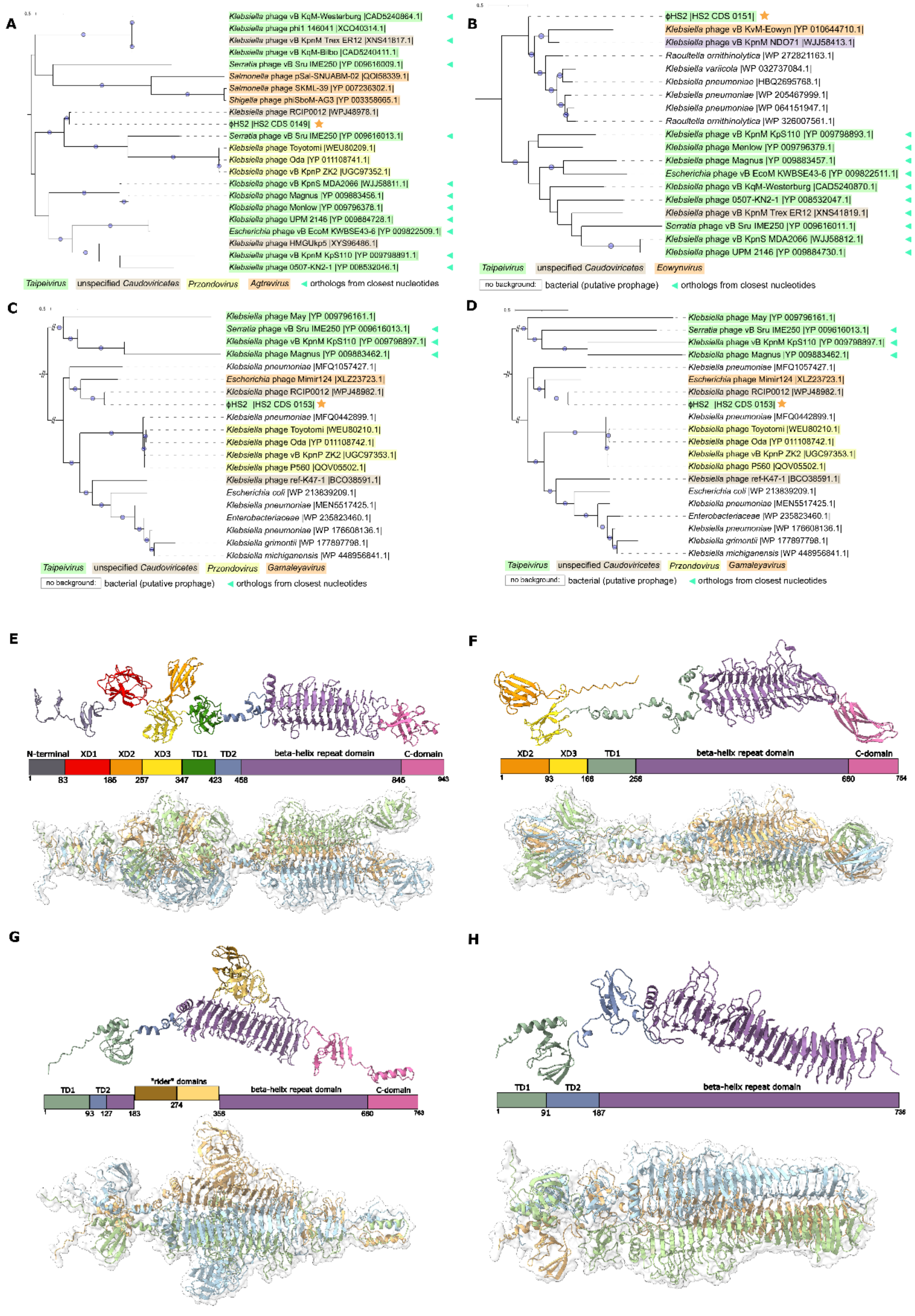


**Figure EV9. Phylogenetic analysis and structural models of ϕHS2 tail spike proteins. (A–D)** Phylogenetic trees of ϕHS2 tail spike protein homologs. Midpoint-rooted maximum likelihood trees are shown for four tail spike proteins: **(A)** CDS_0149, **(B)** CDS_0151,

**(C)** CDS_0153, and **(D)** CDS_0155. Homologs were obtained from the closest nucleotide orthogroup (green triangles) and by BLASTP searches. Background color indicates the lowest taxonomic rank; bacterial protein labels denote the organism group corresponding to the NCBI Identical Protein Groups entry. **(E–H)** AlphaFold3 structural models of ϕHS2 tail spike proteins. Predicted structures of the monomer and homotrimeric complex are shown for **(E)** CDS_0149, **(F)** CDS_0151, **(G)** CDS_0153, and **(H)** CDS_0155. Monomer domains are colour-coded according to the scale bar, with numbers indicating amino acid boundaries. Domain assignments are based on structural representations, HHpred searches and protein alignments, following previously described classifications (Plattner *et al*, 2019; Chao *et al*, 2022).

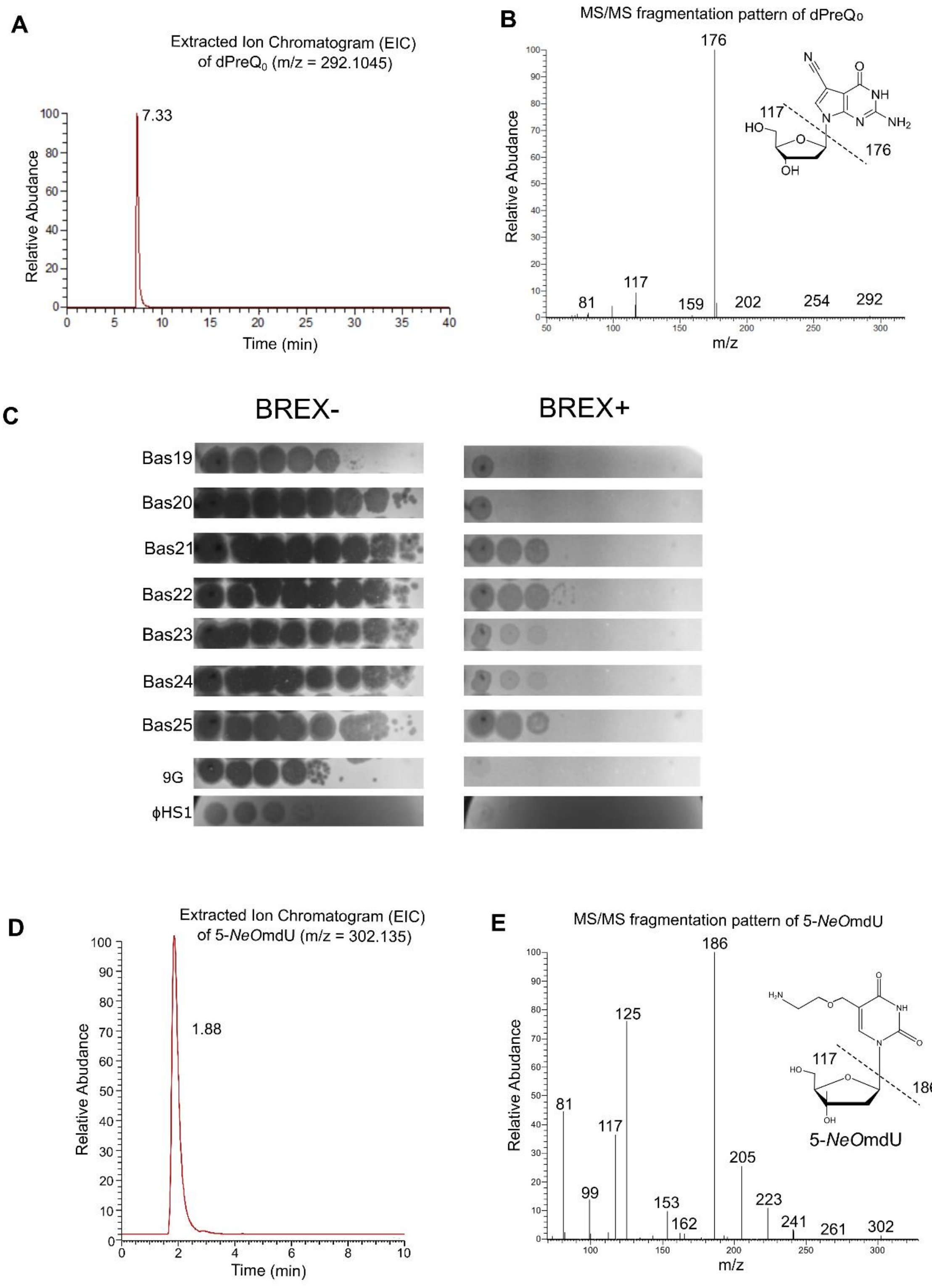


**Figure EV10. Mass spectrometry confirmation of DNA hypermodifications in ϕHS1 and ϕHS2 and assessment of the role of dPreQ$_0$ modification in BREX resistance. (A)** Extracted ion chromatogram (EIC) of dPreQ$_0$ in ϕHS1 genomic DNA. The EIC for the $[M+H]^+$ ion of dPreQ$_0$ (m/z 292.1045) is shown for ϕHS1. A distinct peak (retention time 0–40 min) was detected exclusively in the ϕHS1 sample, confirming the presence of $dPreQ_0$. Abundance is shown in arbitrary units (a.u.). **(B)** MS/MS fragmentation spectrum of $dPreQ_0$ isolated from ϕHS1 genomic DNA. The CID spectrum of the precursor ion $[M+H]^+$ at m/z 292.1 shows a major fragment at m/z 176.1, corresponding to the loss of deoxyribose (117 Da). Subsequent dehydration of the protonated nucleobase yields the ion at m/z 159.1. The proposed fragmentation pathway is indicated. **(C)** Efficiency of plating (EOP) of hypermodified phages 9G and Bas19–25 on BREX-proficient and BREX-deficient hosts. EOP was assessed on *E. coli* BW25113 ($BREX^-$) and BW25113 harbouring the plasmid pBREXAL ($BREX^+$). **(D)** Extracted ion chromatogram (EIC) of 5-NeOmdU in ϕHS2 genomic DNA. The EIC for the $[M+H]^+$ ion of 5-NeOmdU (m/z 302.135) is shown for ϕHS2. A distinct peak was detected exclusively in the ϕHS2 sample, confirming the presence of 5-NeOmdU. Abundance is shown in arbitrary units (a.u.). **(E)** MS/MS fragmentation spectrum of 5-NeOmdU isolated from ϕHS2 genomic DNA. The CID spectrum of the precursor ion $[M+H]^+$ at m/z 302.1 shows a major fragment at m/z 186, corresponding to the protonated nucleobase 5-(2-aminoethoxymethyl)uracil, formed by the loss of protonated 2′-deoxyribose (m/z 117). The fragment at m/z 125 corresponds to protonated 5-hydroxymethyluracil (5-hmU), resulting from the loss of ethanolamine ($HOCH_2CH_2NH_2$) from the m/z 186 ion. Additional fragments derived from the deoxyribose moiety are observed at m/z 81 (dehydrated sugar fragment) and m/z 99 (intermediate dehydration product). The peak at m/z 153 likely originates from a contaminant or background noise and is not diagnostic for 5-NeOmdU.

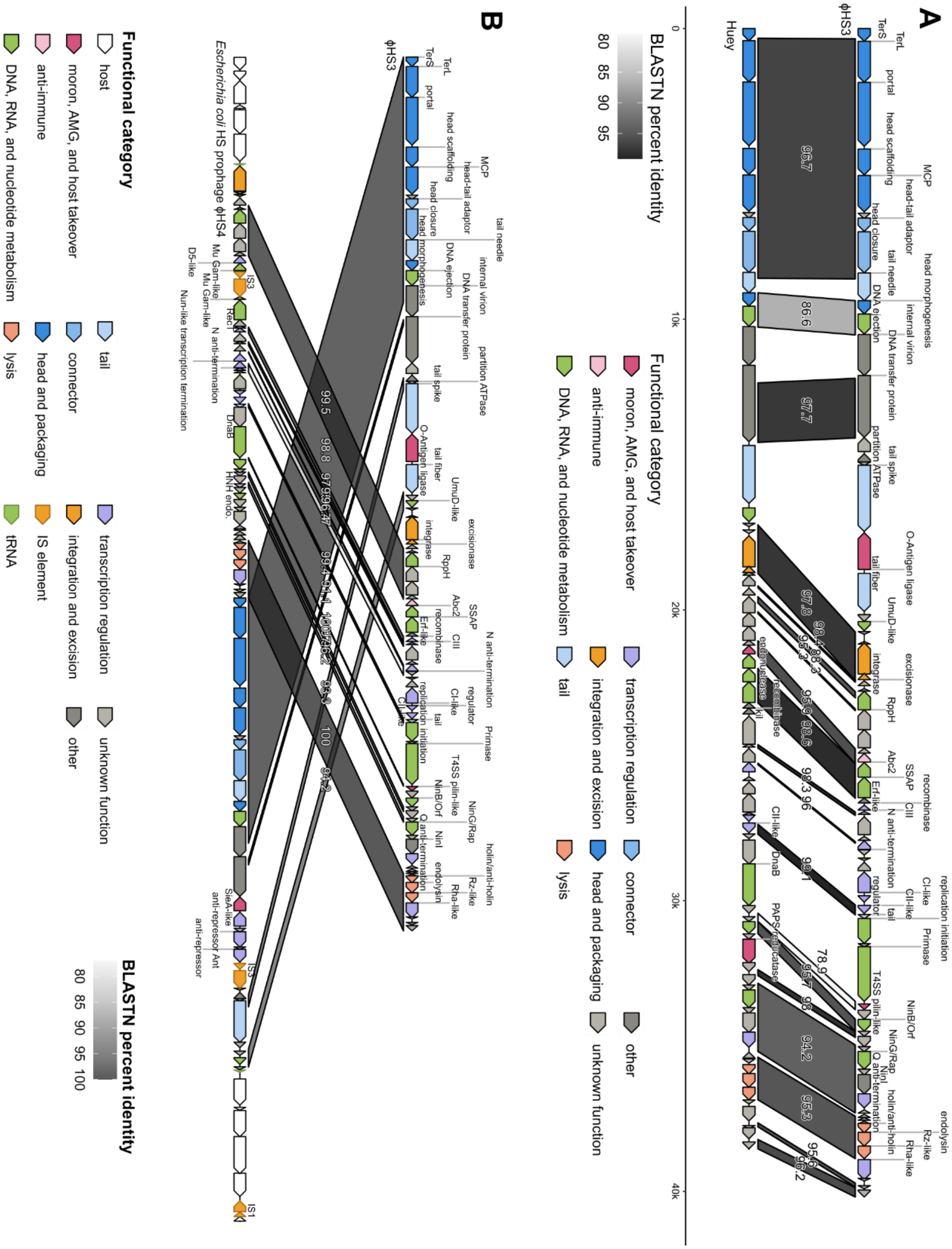


**Figure EV11. Genomic comparison of ϕHS3 with related phages. (A)** Genome organization of *Escherichia* phages ϕHS3 and Huey. Genes are colored according to the predicted function of their products; links indicate regions of DNA sequence similarity. **(B)** Genome organization of *Escherichia* phage ϕHS3 and the *E. coli* HS prophage ϕHS4. Genes are colored according to the predicted function of their products; links indicate regions of DNA sequence similarity.

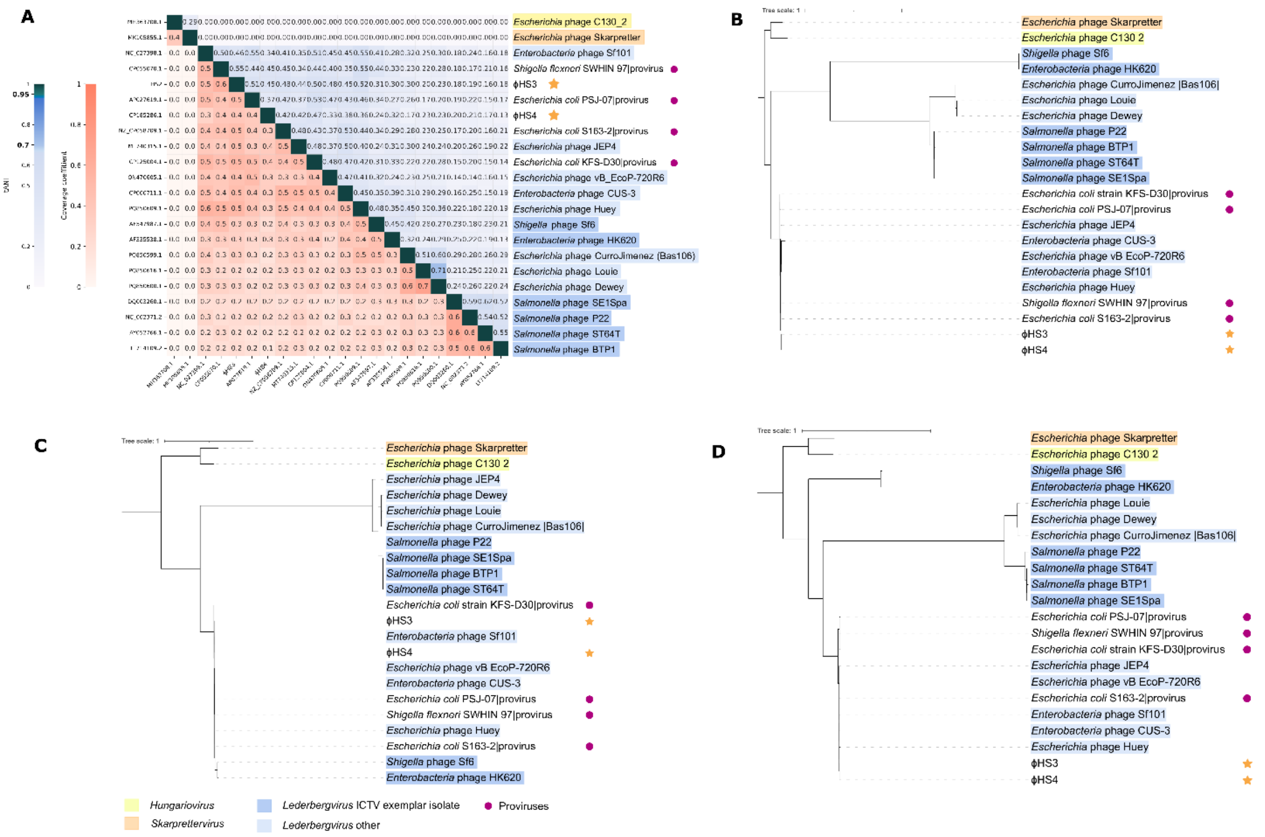


**Figure EV12. Nucleotide identity and phylogenetic analysis of ϕHS3 and related phages. (A)** Total average nucleotide identity (tANI) and coverage matrix constructed using Vclust for the set of closest related genomes, with representatives of the outgroup genera *Skarprettervirus* and *Hungariovirus*. Zero values indicate genome pairs that do not meet the minimum similarity threshold. ICTV-recognized exemplar isolates of the genus *Lederbergvirus* are colored distinctly from those with taxonomy inferred from NCBI GenBank. Genomes labeled "provirus" correspond to prophage regions identified by geNomad in bacterial chromosomes. ϕHS3 and ϕHS4 are indicated by gold stars. **(B)** Midpoint-rooted maximum likelihood phylogenetic tree of the ϕHS3 major capsid protein. **(C)** Midpoint-rooted maximum likelihood phylogenetic tree of the ϕHS3 large terminase subunit. **(D)** Midpoint-rooted maximum likelihood phylogenetic tree of the ϕHS3 portal protein. Trees were constructed using IQ-TREE with the automatically determined best-fit substitution model. Leaf labels are color-coded according to the taxonomic groups shown in the legend.

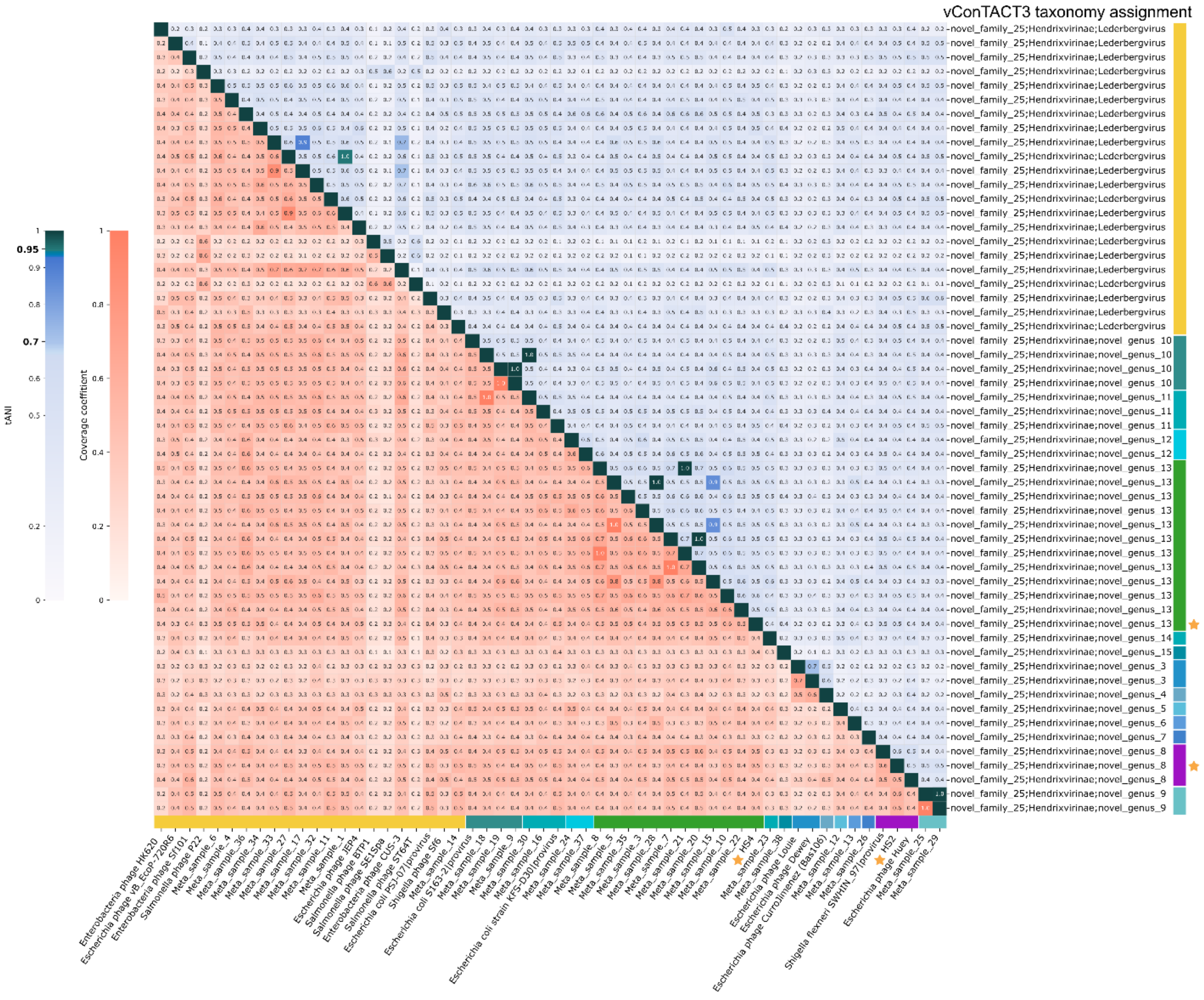


**Figure EV13. Comparison of the closest ϕHS3-related genomes and metavirome contigs.** Total average nucleotide identity and coverage matrix constructed using Vclust. Corresponding taxonomy assignment by vConTACT3 is shown in the right labels. Colored bars indicate different genera (all are proposed as novel except *Lederbergvirus*, marked with a yellow bar). ϕHS3 and ϕHS4 are indicated by gold stars.

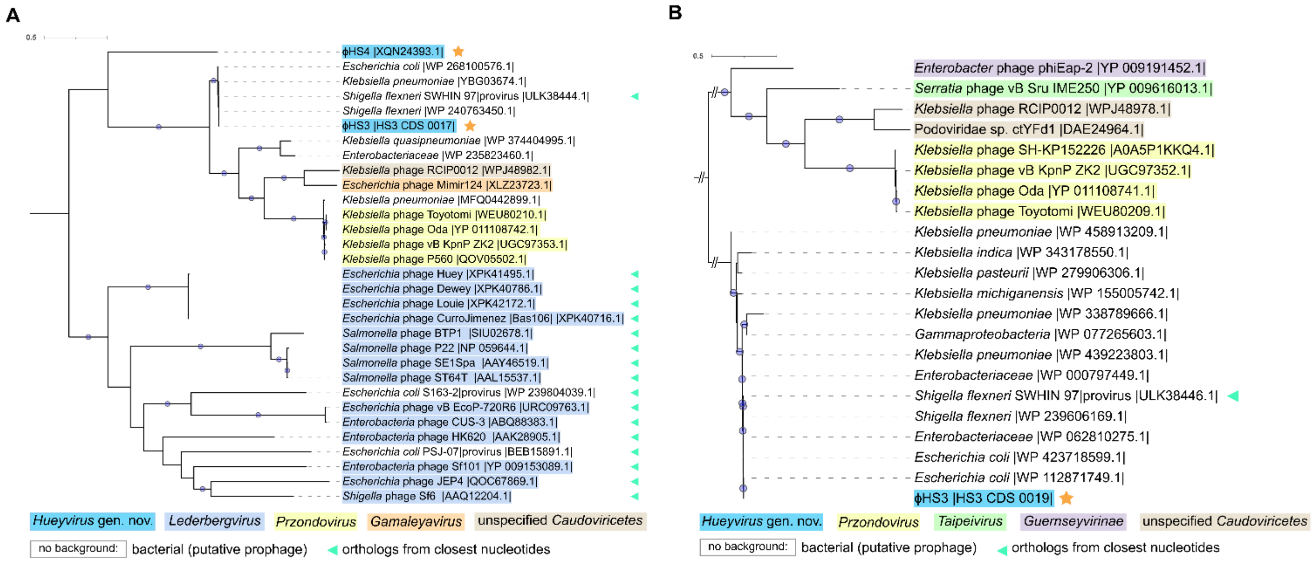


**Figure EV14. Phylogenetic analysis of ϕHS3 tailspike protein homologs**. Midpoint-rooted maximum likelihood phylogenetic trees of ϕHS3 tailspike protein homologs obtained from the closest nucleotide orthogroup (marked with green triangles) and by BLASTP search. Trees are shown for **(A)** CDS_0017 and **(B)** CDS_0019. The background color for phages indicates the lowest taxonomic rank. Bacterial protein labels indicate the organism group corresponding to the NCBI Identical Protein Groups entry.

**Supplementary Table S1**. The reference variants correspond to our assembly.

| Position | Variant type | Variant location | HGVS.c* | HGVS.p* |
|---|---|---|---|---|
| 392133 | missense | 30S ribosomal protein S12 | c.128C>A | p.Thr43Lys |
| 986529 | missense | IS1 family transposase | c.37C>T | p.Pro13Ser |
| 1129338 | synonymous | IS1 family IS1A transposase ORF A | c.207T>A | p.Thr69Thr |
| 1193489 | synonymous | IS1 family IS1A transposase ORF A | c.207T>A | p.Thr69Thr |
| 1244224 | intergenic | IS1 family transposase-IS1 family IS1A transposase ORF A | n.1244224C>T | |
| 1244260 | intergenic | IS1 family transposase-IS1 family IS1A transposase ORF A | n.1244260A>C | |
| 1244442 | synonymous | IS1 family IS1A transposase ORF A | c.207T>A | p.Thr69Thr |
| 1857861 | missense | IS1329 transposase A | c.143C>A | p.Pro48Gln |
| 3149850 | synonymous | RHS repeat protein | c.750G>A | p.Glu250Glu |
| 3149887 | missense | RHS repeat protein | c.713A>C | p.Lys238Thr |
| 3149889 | synonymous | RHS repeat protein | c.711T>C | p.Ser237Ser |
| 3149894 | missense | RHS repeat protein | c.706A>G | p.Ile236Val |
| 3149898 | synonymous | RHS repeat protein | c.702C>G | p.Ala234Ala |
| 3149991 | synonymous | RHS repeat protein | c.609T>G | p.Thr203Thr |
| 3687766 | intergenic | 16S ribosomal RNA–D-glycero-beta-D-manno-heptose-1,7-bisphosphate 7-phosphatase | n.3687766A>G | |
| 4488729 | intergenic | 16S ribosomal RNA–Protoporphyrinogen IX dehydrogenase [quinone] | n.4488729G>T | |

*HGVS.c, HGVS.p refer to the coding DNA and protein levels reference sequence notation used in the Human Genome Variation Society nomenclatu

**Supplementary Table S2. Mutations identified in phage-resistant escape mutants of *E. coli* HS**

| Sample | Position | Reference allele | Alternative allele | Variant type | Gene | Product | HGVS.c_ANN | HGVS.p_ANN |
|---|---|---|---|---|---|---|---|---|
| *E. coli* HS phiHS1 resistant | 4220618 | G | A | missense variant | mdtO | multidrug efflux transporter permease subunit MdtO,UniRef:UniRef50_A0A7D6YZA0 | c.461G>A | p.Arg154His |
| *E. coli* HS phiHS1 resistant | 1756303 | G | T | missense variant | manC | mannose-1-phosphate guanylyltransferase/mannose-6-phosphate isomerase,UniRef:UniRef50_A0A447P8J4 | c.425G>T | p.Gly142Val |
| *E.coli* HS phiHS2 resistant | 864377 | C | T | missense variant | HEAGAH_04385 | NADPH-Fe(3+) oxidoreductase subunit beta,UniRef:UniRef50_A0A6P1Q5Y9 | c.2951G>A | p.Arg984Gln |
| *E.coli* HS phiHS2 resistant | 939596 | C | T | synonymous variant | recB | RecBCD enzyme subunit RecB,UniRef:UniRef50_A0A2X4VHK6 | c.3018C>T | p.Gly1006Gly |
| *E.coli* HS phiHS2 resistant | 1740071 | A | IS1 insert (779 bp) | stop gained & disruptive inframe insertion | wcaA | Putative colanic acid biosynthesis glycosyl transferase WcaA,UniRef:UniRef50_P77414 | c.215A>* | p.Asn70_Gly71ins |

**Supplementary Table S3. Nucleoside peak areas for λ phage control (canonical DNA)**

| Nucleoside | RT (min) | Area | %Area |
|---|---|---|---|
| dC | 1.37 | 634,623 | 14.71 |
| dG | 2.71 | 1,295,467 | 30.02 |
| dT | 4.04 | 876,277 | 20.31 |
| dA | 4.45 | 1,508,516 | 34.96 |

**Supplementary Table S4. Nucleoside peak areas for phage ϕHS1 ($dPreQ_0$-modified DNA)**

| Nucleoside | RT (min) | Area | %Area |
|---|---|---|---|
| dC | 1.37 | 1,161,271 | 13.18 |
| dG | 2.72 | 1,736,958 | 19.72 |
| dT | 4 | 1,980,336 | 22.48 |
| dA | 4.46 | 3,443,135 | 39.08 |
| $dPreQ_0$ | 7.13 | 488,243 | 5.54 |

**Supplementary Table S5. Nucleoside peak areas for phage ϕHS2 (5-NeOmdU-modified DNA)**

| Nucleoside | RT (min) | Area | %Area |
|---|---|---|---|
| dC | 1.19 | 1450898.054 | 15.52 |
| 5-NeOmdU | 1.43 | 427400.91 | 4.57 |
| dG | 2.87 | 2573574.77 | 27.54 |
| dT | 4.18 | 714653.5 | 7.65 |
| dA | 4.46 | 3423186.973 | 36.63 |

**Supplementary Table S7. Bacterial strains, plasmids, and phages used in the study.**

| *Strains* | **Comments** | **Source** |
|---|---|---|
| *E.coli* HS | Natural isolate with $BREX^{Ec}$, *$Str^R$* | Paul Cohen |
| *E.coli* BW25113 *E.coli* | Strain for routine work | Lab stock |
| *E.coli* HS ϕHS3 lysogen | HS strain with lysogenized ϕHS3 phage | This work |
| *E.coli* HSΔP22 (ϕHS4) | HS strain with a deletion of the P22 prophage | This work |
| *E.coli* HS *brxX(Q23*)* | HS strain with impaired BrxX (PglX) methyltransferase production | This work |
| *E.coli* HS *wbdC(Q181*)* | HS strain with impaired O-antigen synthesis | This work |
| *E.coli* ECOR-collection | Collection resembling a variety in the O-antigen structures (72 strains) | (Ochman & Selander, 1984) |
| *K. pneumonia* B-9185 (K47) | | State Collection of Pathogenic Microorganisms and Cell Cultures «SCPM-Obolensk» |

| | | |
|---|---|---|
| *K. pneumoniae* (capsule type K1, K2, K10, K20, K23, K39, K57) | | Lab stock |
| **Phages** | **Comments** | **Source** |
| ϕHS1-3 | | This work |
| BASEL collection | BASEL collection (1-106 phages) | (Maffei *et al*, 2021; Humolli *et al*, 2025) |
| **Plasmids** | | |
| pScI_dCas–CDA_J23119–sgRNA | Bacterial Target-AID vector with sgRNA (dCas9-CDA base editor for stop codon introduction) | (Banno *et al*, 2018) |
| pBREX AL | 6-genes *E. coli* HS BREX cluster cloned under native promoter in pBTB-2, KanR | (Gordeeva *et al*, 2019) |

**Supplementary Table S8. Primers used in the study.**

| Name | Sequence | Purpose |
|---|---|---|
| brxX_stop_spacer_F | TAGCTGGGAGCAGGTAGAGGTTAT | Cloning of *pglX*-targeting spacer for stop-knockout (dCas9-CDA) |
| brxX_stop_spacer_R | AAACATAACCTCTACCTGCTCCCA | (reverse complement sequence) |
| brxX_seq_F | ACTGCTGCTGCCGGATAAC | Sanger sequencing of *brxX* to verify stop codon introduction |
| brxX_seq_R | GCTCAATGTAGTACTCCATTTTG | Sanger sequencing of *brxX* (reverse direction) |
| wbdC_stop_spacer_F | TAGCCTCAATACCGCCGTAGGTAT | Cloning of *wbdC*-targeting spacer for stop-knockout (dCas9-CDA) |
| wbdC_stop_spacer_R | AAACATACCTACGGCGGTATTGAG | (reverse complement sequence) |
| wbdC_seq_F | GTAGGCGACAGGACCTGTC | Sanger sequencing of *wbdC* to verify stop codon introduction |
| wbdC_seq_R | GTCACTTCTTTGCCTGAGGTG | Sanger sequencing of *wbdC* (reverse direction) |
| ϕHS4_qPCR_F | TGGAGATAGTGACTGGTACTG | qPCR quantification of P22-like prophage (ϕHS4) genome amplification after mitomycin C induction |
| ϕHS4_qPCR_R | CCAATGTATCCTAATATTCTTGTTAG | qPCR quantification of P22-like prophage (ϕHS4) genome amplification after mitomycin C induction |
| DuoHS_qPCR_F | CTTTCCGCGAGTCGGTATTG | qPCR quantification of P2-like prophage (DuoHS) genome amplification after mitomycin C induction |

| | | |
|---|---|---|
| DuoHS_qPCR_R | TTTGGCTTCAGGGAGATATCC | qPCR quantification of P2-like prophage (DuoHS) genome amplification after mitomycin C induction |
| ϕHS3_check_F | TGCGCAATCATTGGTGTTGG | Detection of ϕHS3 lysogen |
| ϕHS3_check_R | AATGCAGTTTATAGCCCCCTCG | Detection of ϕHS3 lysogen |
| gyrA_F1 | CGTACTTTACGCCATGAACGTACTAGGC | Real-time PCR of *gyrA* gene |
| gyrA_R1 | GCCATGCGGACGATCGTGTCATAG | Real-time PCR of *gyrA* gene |
| ϕHS4 attB-flank F | GTGGCAAAGAAACGTGTCGCAC | used to detect P22-like (ϕHS4) prophage excision by PCR |
| ϕHS4 attB-flank R | TTCATACGCTCGGCAGGTATCTC | used to detect P22-like (ϕHS4) prophage excision by PCR (reverse direction) |
| P22-F | CTTTCCGCGAGTCGGTATTG | Primers for detection of P22-like (ϕHS4) prophage |
| P22-R | TTTGGCTTCAGGGAGATATCC | Primers for detection of P22-like (ϕHS4) prophage (reverse direction) |
| ϕHS3 attB-flank F | CAGGCGCGGTTTGATCAG | Integration site of ϕHS3 prophage in *E. coli* HS genome |
| ϕHS3 attB-flank R | TTTAACGTTCATTTCCACTCTCTGG | Integration site of ϕHS3 prophage in *E. coli* HS genome (reverse direction) |
| ϕHS3 attL R | CGCTAATGCTCTGTTACAGGTC | attL site of ϕHS3 |
